# XIST dampens X chromosome activity in a SPEN-dependent manner during early human development

**DOI:** 10.1101/2023.10.19.563078

**Authors:** Charbel Alfeghaly, Gaël Castel, Emmanuel Cazottes, Madeleine Moscatelli, Eva Moinard, Miguel Casanova, Juliette Boni, Kasturi Mahadik, Jenna Lammers, Thomas Freour, Louis Chauviere, Carla Piqueras, Ruben Boers, Joachim Boers, Joost Gribnau, Laurent David, Jean-François Ouimette, Claire Rougeulle

**Affiliations:** Université Paris Cité, CNRS, Epigenetics and Cell Fate, F-75013 Paris, France; Nantes Université, CHU Nantes, Inserm, CR2TI, F-44000 Nantes, France; Department of Developmental Biology, Erasmus University Medical Center Rotterdam, Rotterdam, Netherlands; CHU Nantes, Service de Biologie de la Reproduction, Nantes, France; Nantes Université, CHU Nantes, Inserm, CNRS, BioCore, F-44000 Nantes, France

## Abstract

XIST long non-coding RNA is responsible for X chromosome inactivation (XCI) in placental mammals, yet it accumulates on both X chromosomes in human female pre-implantation embryos without triggering X chromosome silencing. The long non-coding RNA XACT co-accumulates with XIST on active Xs and may antagonize XIST function. Here we used human ES cells in a naïve state of pluripotency to assess the function of XIST and XACT in shaping the X chromosome chromatin and transcriptional landscapes during pre-implantation development. We show that XIST triggers the deposition of polycomb-mediated repressive histone modifications and attenuates transcription of most X-linked genes in a SPEN-dependent manner, while XACT deficiency does not significantly affect XIST activity or X-linked gene expression. Our study demonstrates that XIST is functional prior to XCI, confirms the existence of a transient process of X chromosome dosage compensation, and reveals that X chromosome inactivation and dampening rely on the same set of factors.

## Introduction

In mammals, males and females differ genetically by the presence of sex chromosomes, with females carrying 2 X chromosomes and males one X and a Y chromosome. The human X chromosome is one of the seven largest chromosomes, harboring ∼1100 genes and 4% of the total number of protein-coding genes, both housekeeping and tissue-specific. X chromosome dosage imbalance between males and females is compensated by the transcriptional inactivation of one of the 2 Xs in females (Avner and Heard, 2001; Brockdorff et al., 2020; Plath et al., 2002). In the mouse, X chromosome inactivation (XCI) is set up during early female embryogenesis, soon after zygotic genome activation (Hadjantonakis et al., 2001; Sugimoto et al., 2000; Tan et al., 1993). Such a timing suggests that functional compensation of X chromosome dosage is essential not only for cellular function and homeostasis in adults, but also for early development, a hypothesis that is confirmed by the female embryonic lethality caused by improper XCI establishment (Marahrens et al., 1997; Penny et al., 1996).

In contrast to rodents, primate female pre-implantation development proceeds in the absence of XCI (Okamoto et al., 2021, 2011). Strikingly, XIST, which is the trigger of XCI, is already expressed at these early stages in humans and XIST RNA decorates both active X chromosomes (Okamoto et al., 2011; Vallot et al., 2017). XIST RNA has been shown in the mouse to recruit a plethora of protein factors that induce a cascade of remodeling events leading to transcriptional silencing and chromatin reorganization of the inactivating X chromosome (Chu et al., 2015; McHugh et al., 2015). One of these effectors is SPEN, which plays a key role in the initiation and establishment of XCI (Dossin et al., 2020; Żylicz et al., 2019). SPEN binds XIST *via* its RNA recognition motifs (RRM) and interacts, *via* its SPOC domain, with multiple factors, including the corepressors NCoR/SMRT and the NuRD complex, to activate histone deacetylase 3 (HDAC3) and to promote transcriptional repression (Dossin et al., 2020; Żylicz et al., 2019). In addition, XIST indirectly recruits the noncanonical polycomb repressive complex (PRC) 1, leading to the accumulation of repressive histone modifications such as H2AK119Ub and to PRC2-mediated H3K27me3, which are hallmarks of the inactive X (Bousard et al., 2019; Nakamoto et al., 2020; Pintacuda et al., 2017a). PRC1 and SPEN have been shown to act in parallel in the repression of X-linked genes, while PRC2 appears mostly dispensable (Bowness et al., 2022).

The uncoupling between XIST expression and XCI establishment in human early development raises questions as to whether XIST is functional at this stage. XIST ability to recruit enzymatically active histone modifying complexes is challenged by the reported lack of accumulation of repressive histone modifications on XIST-coated X chromosomes in human pre-implantation embryos (Okamoto et al., 2011). Additionally, whether XIST intervenes in other transcriptional compensation mechanisms which may operate at this specific time window, is an open question. Single-cell RNA-seq (scRNA-seq) of human pre-implantation embryos indeed revealed a progressive decrease, during the first days of female embryogenesis, of X-linked gene expression from both active X chromosomes, a process referred to as X chromosome dampening (XCD) (Petropoulos et al., 2016). Finally, the presence of yet unknown factors preventing XIST from silencing the X chromosome remains to be determined.

We have previously identified XACT, a primate specific lncRNA that is expressed and accumulates *in cis* on both active X chromosomes during early human development (Casanova et al., 2019; Vallot et al., 2017, 2013). The expression of XACT is mainly restricted to pluripotent stages, suggesting a role for XACT in this cellular context (Vallot et al., 2013). Previous studies have shown that XIST and XACT RNAs occupy distinct nuclear territories in human pre-implantation embryos and that XACT expression in transgenic mouse cells influences XIST accumulation *in cis* (Vallot et al., 2017). This renders XACT an interesting candidate which might act as an XCI antagonist by affecting either XIST expression, localization or activity.

Here we capitalized on female human embryonic stem cells (hESCs), which can be transitioned from primed to naïve state of pluripotency capturing post- and pre-XCI status, respectively, to study X chromosome activity and dosage compensation mechanisms in early human development. Using constitutive and inducible loss-of-function (LOF) approaches, we demonstrate that XACT is not impacting X-linked gene expression in naïve hESCs, despite XIST being functional in this context. Indeed, we show that XIST is responsible for a reduction in most, but not all X-linked gene transcription levels and for the accumulation of PRC-mediated repressive histone marks on active X chromosomes. Yet, we uncover massive redistribution of XIST RNA and PRC-mediated histone modifications on dampened compared to inactive X chromosomes. We also report enrichment in H3K27me3 on X chromosomes in embryos, thus resolving a long-standing question in the field and reconciling *in vitro* models to *ex vivo* embryos. Finally, we provide evidence that XIST interacts with SPEN prior to XCI, and reveal a direct role of SPEN in X chromosome dampening. Overall, our study demonstrates that a core set of factors triggers distinct dosage compensation mechanisms in a developmental stage dependent manner.

## Results

### XIST accumulation attenuates X chromosome transcription in naïve hESCs

To study the X chromosome status in early development and to probe the function of XIST prior to XCI establishment, we took advantage of primed to naïve reprogramming of H9 hESCs using NaïveCult (Stemcell™ technologies) and PXGL media (Bredenkamp et al., 2019; Guo et al., 2017). We validated through transcriptomic analyses the identity of reset cells, which expressed naïve-specific pluripotency markers (Figure 1A) and display transcriptomic signatures of pre-implantation epiblasts (Petropoulos et al., 2016; Zhou et al., 2019), while primed H9 hESCs resemble the post-implantation epiblast (Figure 1B). We also verified the X chromosome activity status using a list of X-linked informative single nucleotide polymorphisms (SNPs) distributed over at least 140 genes (Supplementary Table 1). This confirmed that X chromosome reactivation occurred, with biallelic expression of the vast majority of tested X-linked SNPs (87% and 75% in NaïveCult and PXGL naïve hESCs, respectively, compared to 12% in primed hESCs (Figure 1C and Supplementary Figure 1A). X chromosome reactivation was confirmed at the single cell level using RNA fluorescent *in situ* hybridization (FISH) (Figure 1D and Supplementary Figure 1B). Concordantly, two XACT clouds were also detected in 97% of naïve cells as previously described in human pre-implantation embryos (Vallot et al., 2017) (Figure 1E and Supplementary Figure 1B). Consistent with previous reports (Sahakyan et al., 2017; Vallot et al., 2017), XIST was expressed and mostly accumulated on one of the two X chromosomes in both NaïveCult and PXGL naïve cells (Figure 1D, E and Supplementary Figure 1B). Allelic analysis shows that XIST expression is mainly from the same X chromosome in naïve and primed hESCs (Supplementary Figure 1C).

**Figure 1:**
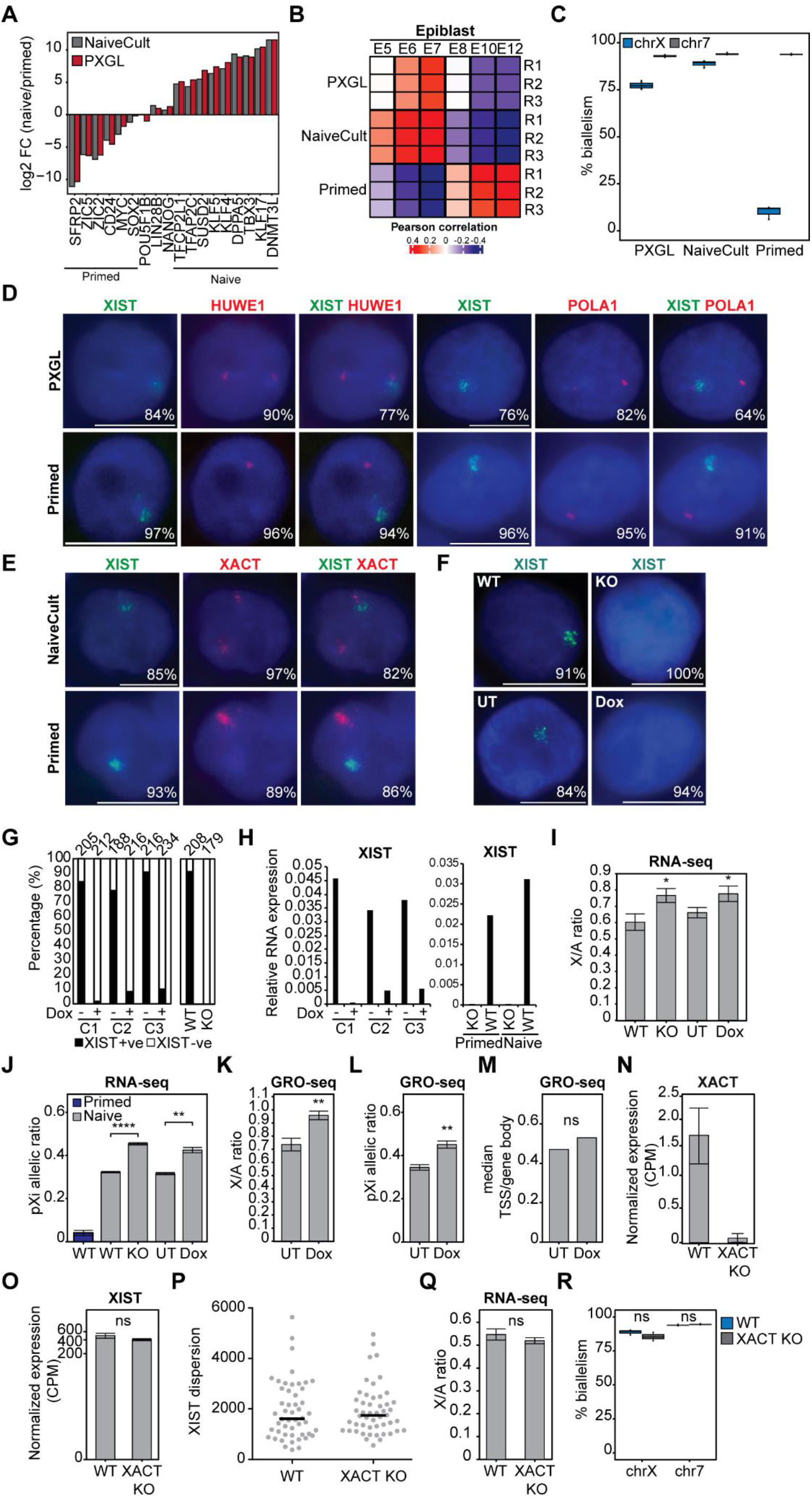
XIST accumulation attenuates X chromosome transcription in naïve hESCs. **A)** Bar plot showing log2 FC between naïve (NaïveCult and PXGL) and primed H9 hESCs obtained from RNA-seq data. Core pluripotency factors as well as markers of each pluripotent state are highlighted. **B)** Correlation heatmap generated using epiblast markers specific to each blastocyst stage from human pre-implantation and post-implantation embryo scRNA-seq data (Petropoulos et al., 2016; Zhou et al., 2019). Numbers correspond to the embryonic days (E5 to E12). **C)** Percentage of biallelically expressed SNPs from X chromosome and chromosome 7 in naïve (NaiveCult and PXGL) and primed H9 hESCs. **D)** Representative RNA-FISH images for XIST and two X-linked genes (HUWE1 and POLA1) in naïve (PXGL) and primed H9 hESCs. Percentages of cells displaying the representative pattern are indicated. Scale bar= 10 µm. **E)** Representative RNA-FISH images for XIST and XACT in naïve (NaiveCult) and primed H9 hESCs. Percentages of cells displaying the representative pattern are indicated. Scale bar= 10 µm. **F)** Representative RNA-FISH images for XIST in naïve H9 hESCs with a stable or doxycycline inducible XIST repression. Percentages of cells displaying the representative pattern are indicated. Scale bar= 10 µm. **G)** Quantification of RNA-FISH data showing the percentage of cells with XIST signal in XIST WT, KO and in 3 XIST CRISPRi naïve H9 clones (C1-C3) treated or not with doxycycline. **H)** RTqPCR analysis of XIST expression in naïve XIST CRISPRi clones with or without doxycycline treatment (left), and in WT and XIST KO naïve and primed H9 hESCs (right). Values are normalized to β-actin. **I)** X/A ratio from RNA-seq data of WT, XIST KO, XIST CRISPRi untreated (UT) and Dox-treated (Dox) naïve H9 cells (n=3). Error bars correspond to the standard deviation (SD) of replicates. **J)** pXi allelic ratio from RNA-seq data of primed (WT) and naïve (WT, XIST KO, XIST CRISPRi UT and Dox-treated) H9 hESCs (n=3). Error bars correspond to the SD of replicates. **K)** X/A ratio from fastGRO-seq data of XIST CRISPRi UT and Dox-treated naïve H9 cells (n=3). Error bars correspond to the SD of replicates. **L)** pXi allelic ratio from fastGRO-seq data of XIST CRISPRi UT and Dox-treated naïve H9 cells (n=3). Error bars correspond to the SD of replicates. **M)** Ratio of the median coverage of reads on TSS/gene body from fastGRO-seq data from XIST CRISPRi UT and Dox-treated H9 naïve hESCs (n=3). **N)** XACT expression level in WT and XACT KO naïve hESCs obtained from RNA-seq data (n=3). Error bars correspond to the SD of replicates. **O)** XIST expression level in WT and XACT KO naïve hESCs obtained from RNA-seq data (n=3). Error bars correspond to the SD of replicates. **P)** Distribution of the dispersion of XIST RNA-FISH signal in WT and XACT KO naïve cells. Horizontal line represents the mean dispersion of each group. **Q)** X/A ratio from RNA-seq data of WT and XACT KO naïve H9 cells (n=3). Error bars correspond to the SD of replicates. **R)** Percentage of biallelically expressed SNPs from the X chromosome and chromosome 7 in WT and XACT KO naïve H9 hESCs. Unless otherwise specified, p-values were calculated by two-sided unpaired t-test. Levels of significance: ns≥0.05; * < 0.05; ** < 0.01; *** < 0.001; **** < 0.0001.

We generated XIST loss-of-function (LOF) systems based on (i) CRISPR/Cas9-mediated genome editing and (ii) a doxycycline inducible CRISPR/dCas9 strategy, to deplete XIST expression in H9 hESCs in a constitutive or inducible manner, respectively. One stable homozygous XIST KO (Supplementary Figure 1D) and 3 independent XIST CRISPRi primed hESC clones (Supplementary Figure 1E) were obtained and converted to the naïve state using the PXGL culture regimen.

XIST was efficiently repressed in naïve XIST KO and dox-treated CRISPRi clones, as measured by RNA-FISH and RTqPCR (Figure 1F-H). Of note, the ability to derive XIST KO naïve H9 cells with normal expression of naïve specific and core pluripotency factors indicates that XIST is not required for primed to naïve resetting (Supplementary Figure 1F). Principle Component Analysis (PCA) of RNA-seq data generated from stable XIST KO and inducible XIST CRISPRi clones revealed that naïve XIST +ve (WT and untreated CRISPRi clones) and XIST -ve (KO and Dox-treated CRISPRi clones) hESCs were grouped together and clustered away from primed hESCs, indicating that the principal variation is due to the cellular state and that loss of XIST does not alter the global naïve transcriptome (Supplementary Figure 1G).

We probed the impact of XIST loss on X chromosome activity in naïve hESCs by measuring the X/A ratio, which was around 1.3-fold higher in XIST KO and CRISPRi Dox clones compared to WT and CRISPRi UT clones (Figure 1I). This increase was not due to changes in global autosomal gene expression levels, which remained constant (Supplementary Figure 1H). This indicates a global upregulation of the X chromosome expression in the absence of XIST, which also translates in a slight increase in the percentage of biallelically detected X-linked SNPs in naïve XIST -ve cells compared to XIST +ve cells (Supplementary Figure 1I).

The level of increase (1.3-fold) is compatible with XIST being expressed from and regulating only one X chromosome. This suggests that loss of XIST leads to higher expression from the former XIST-coated X chromosome in naïve hESCs.

We took advantage of the fact that XIST is expressed only from one of the two active X chromosomes in naïve hESCs (the previous Xi, Supplementary Figure 1C) to confirm that the underlying effects are directly linked to the loss of XIST. As all primed H9 hESCs have the same inactive X, we could split, using SNPs, the reads between the X chromosomes, assign them to either the previous Xi (pXi) or previous Xa (pXa) in naïve hESCs, and calculate a pXi allelic ratio. As expected, this ratio was of 0.04 in primed hESCs, indicating that most reads were assigned to the active X chromosome (Figure 1J). In contrast, WT and XIST CRISPRi UT naïve hESCs displayed a pXi allelic ratio of 0.35, indicating that both X chromosomes were active, yet the pXi was less expressed compared to the pXa (Figure 1J). The pXi allelic ratio was significantly increased in the absence of XIST to reach 0.5, indicating equal expression from both X chromosomes (Figure 1J). This demonstrates that XIST accumulation attenuates X chromosome expression in *cis* in naïve hESCs.

To probe whether XIST-mediated dampening impacts X-linked gene expression at the transcriptional or post-transcriptional level, we measured nascent transcription using the low input fastGRO-seq approach (Barbieri et al., 2020). We observed an increase in the X/A and pXi allelic ratios in the absence of XIST similar to that observed by bulk RNA-seq (Figure 1K, L). This shows that XIST regulates ongoing transcription of X-linked genes in naïve hESCs. As we found no significant difference in the promoter-proximal pausing ratio between untreated and Dox-treated CRISPRi XIST clones (Figure 1M), we concluded that XIST-mediated attenuation acts at the level of transcription initiation. Altogether, our data show that XIST accumulation attenuates X chromosome transcription in naïve hESCs.

### XACT does not control X chromosome activity in naïve hESCs

XIST capacity to attenuate X chromosome transcription instead of triggering complete inactivation made us question whether XACT antagonizes XIST silencing activity. We thus deleted a 90 kb region encompassing XACT transcription start sites (TSS) in primed H9 cells (Figure 1N and Supplementary Figure 1J, N). Three independent homozygous *XACT* KO clones were successfully converted to a naïve pluripotent state. Neither expression of naïve specific and core pluripotency factors, nor cell morphology and growth were impacted by XACT deletion (Supplementary Figure 1K). Furthermore, PCA reveals that XACT KO and WT naïve H9 cells were grouped together and clustered away from primed hESCs, indicating that loss of XACT does not significantly alter the naïve hESC transcriptome (Supplementary Figure 1L). This was confirmed through differential expression analysis, which revealed few genes with altered expression levels (14 upregulated and 16 downregulated; log2FC > |1| and FDR<0.05) in XACT KO compared to WT H9 naïve cells (Supplementary Figure 1M). Gene ontology analysis did not reveal any enriched pathways in the list of the 30 differentially expressed genes (DEGs). This indicates that XACT is dispensable for primed to naïve resetting and plays no major role in naïve hESCs.

XIST RNA levels were not altered in XACT KO clones (Figure 1O). Similarly, the distribution of the XIST signal, which is more dispersed in naïve hESCs compared to primed hESCs and to fibroblasts (Sahakyan et al., 2017; Vallot et al., 2017), is similar in WT and XACT KO clones (Figure 1P and Supplementary Figure 1N), indicating that XACT does not regulate XIST expression or localization in naïve hESCs. As a proxy for global X chromosome expression, we measured the X/A ratio and observed no significant changes in naïve XACT KO compared to WT hESCs (Figure 1Q). Furthermore, the percentage of biallelic X-linked SNPs detected by RNA-seq in XACT KO cells remained unchanged compared to WT, indicating that X chromosome silencing did not initiate in the absence of XACT in naïve hESCs (Figure 1R). These results indicate that XACT is not an antagonist of XIST and does not control X chromosome activity in naïve hESCs.

### XIST is broadly distributed on the dampened X chromosome in naïve hESCs

The pattern of XIST accumulation in human pre-implantation embryos and naïve hESCs is thought to differs from that of primed hESCs, being more dispersed as shown by RNA-FISH (Sahakyan et al., 2017; Vallot et al., 2017). To address whether change in XIST repressive activity between naïve and primed pluripotency could be linked to redistribution of XIST RNA, we mapped at high resolution the genomic binding sites of XIST on dampened and inactive X chromosomes, in naive and primed H9 hESCs, respectively, using RNA Antisense Purification followed by high-throughput sequencing (RAP-seq) (Engreitz et al., 2013). XIST was efficiently and specifically captured from the chromatin of both cell types (Figure 2A).

**Figure 2:**
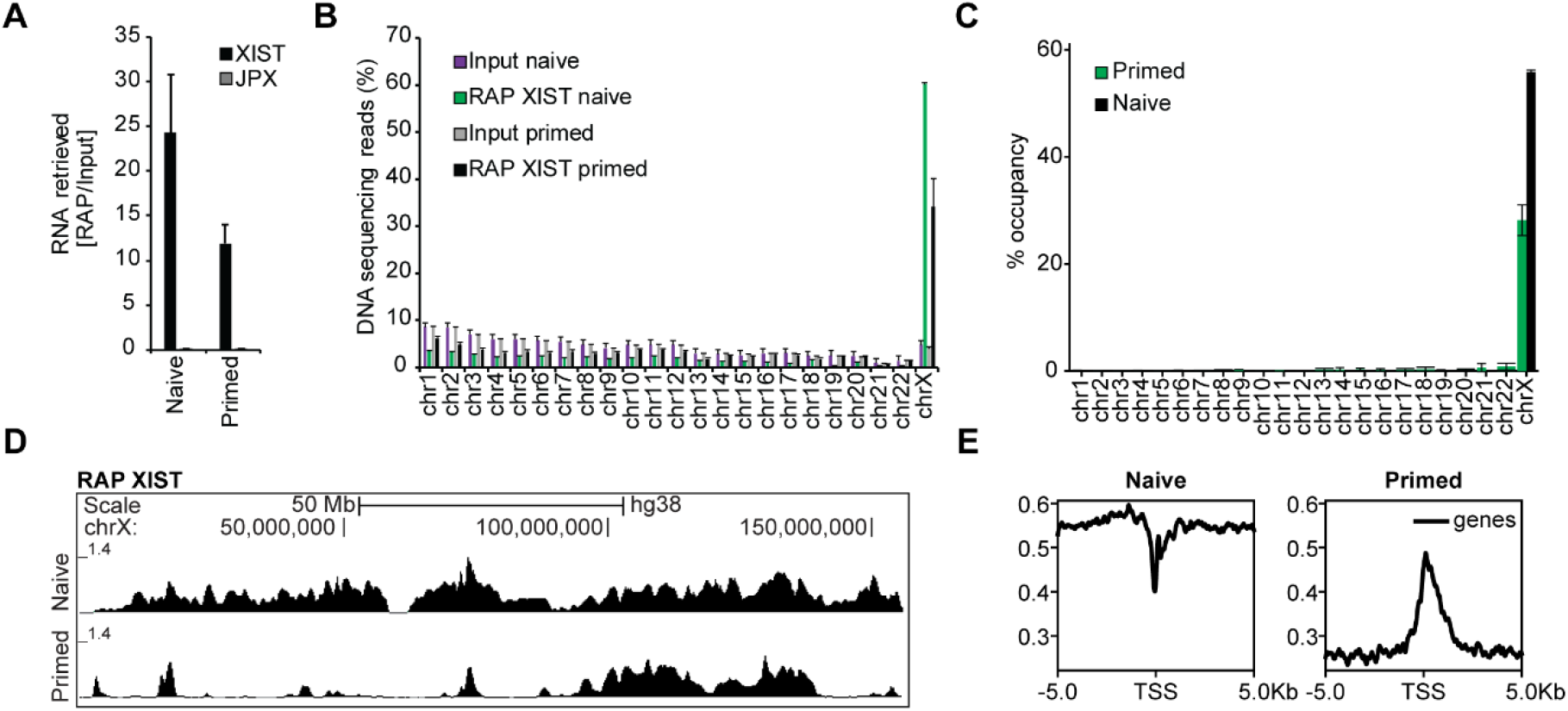
XIST is broadly distributed on the dampened X chromosome in naïve hESCs. **A)** Analysis by RTqPCR of XIST and JPX enrichment after XIST pulldown in naïve and primed H9 hESCs. JPX was used as a negative control. Values were normalized to Input (n=2). Error bars correspond to the SD of replicates. **B)** Percentage of aligned DNA sequencing reads per chromosome after XIST pulldown and in input material in naïve and primed H9 hESCs (n=2). Error bars correspond to the SD of replicates. **C)** Percentage of XIST RAP peak occupancy per chromosome for primed and naïve H9 hESCs (n=2). Error bars correspond to the SD of replicates. **D)** XIST RAP profiles (log2 enrichment over Input) on the X chromosome in naïve and primed H9 hESCs. **E)** Profile plots of XIST RAP-seq reads over 10 kb region flanking TSS of X-linked genes in naïve and primed H9 hESCs.

The X chromosome showed the highest percentage of mapped DNA sequencing reads in both naïve (60%) and primed (34%) hESCs, while the remaining reads were distributed across all autosomes (Figure 2B). Principle component and matrix correlation analyses show that the naïve RAP XIST replicates cluster together and away from the primed RAP XIST replicates. As the comparison is made in an isogenic framework, we can conclude that variation in XIST distribution is due to the cellular context (Supplementary Figure 2A, B). Indeed, XIST was broadly distributed on the X chromosome in naïve hESCs, occupying >50% of the chromosome, while XIST contacted a reduced fraction (<30%) of the primed X chromosome, mostly on the long arm (Figure 2C, D). Differential enrichment analysis confirmed that most of the X chromosome is enriched for XIST in naïve compared to primed hESCs (Supplementary Figure 2C). However, XIST was overall depleted from X-linked TSS compared to flanking sequences in naïve hESCs. This pattern is in sharp contrast to that of primed hESCs, where XIST accumulated over TSS (Figure 2E). These observations support a role for XIST distribution in the regulation of X-linked genes, where XIST RNA molecules relocate from X-linked promoters to flanking regions upon primed to naïve transition.

We then probed the correlation of XIST distribution with several genomic features in both naïve and primed hESCs. In naïve hESCs, XIST coverage was moderately correlated with density of transposable elements such as LINE and retrotransposons, in contrast to primed hESCs (Supplementary Figure 2D). This could reveal a role for these elements in XIST spreading upon initial XIST up-regulation, as previously suggested (Bailey et al., 2000; Cantrell et al., 2009). Difference in XIST distribution was however not linked to differential XIST expression levels (Supplementary Figure 2E), suggesting that broader coverage of the X chromosome in naïve hESCs is not due to increased number of XIST RNA molecules. XIST transcript reconstruction revealed the presence of multiple XIST isoforms (Supplementary Figure 2F). XIST 1.1 isoform, with 6 exons, corresponds to the annotated XIST structure and is expressed in naïve and primed hESCs. Other isoforms are specific to the primed or naïve contexts and differ in their first and last exon. Importantly, all the isoforms are produced from the same TSS and harbor all 6 repeat elements (A-F) that are important for XIST function and localization (Supplementary Figure 2F) (Pintacuda et al., 2017b). Altogether, these data reveal that XIST establishes distinct contacts on dampened and inactive X chromosomes, in line with distinct activities of XIST in naïve and primed pluripotent contexts.

### Repressive histone modifications accumulate on dampened X chromosome in a XIST-dependent manner in naïve hESCs

Since XIST is known to indirectly recruits PRC complexes, and given its ability to control X chromosome activity in naïve hESCs, we interrogated the chromatin landscape of dampened X chromosomes and its link to XIST. We used CUT&RUN, which provided the opportunity to precisely compare H3K27me3 and H2AK119Ub distributions to that of XIST along X chromosomes in naïve hESCs, and also to that of primed hESCs. We also analyzed H3K9me3, which is enriched on the Xi independently of XIST. Levels of H3K27me3, H2AK119Ub and, to a lesser extent, H3K9me3, were higher on the X chromosomes than on autosomes in naïve hESCs (Figure 3A), even if this tendency was less pronounced than in primed hESCs, at least for H3K27me3 and H3K9me3.

**Figure 3:**
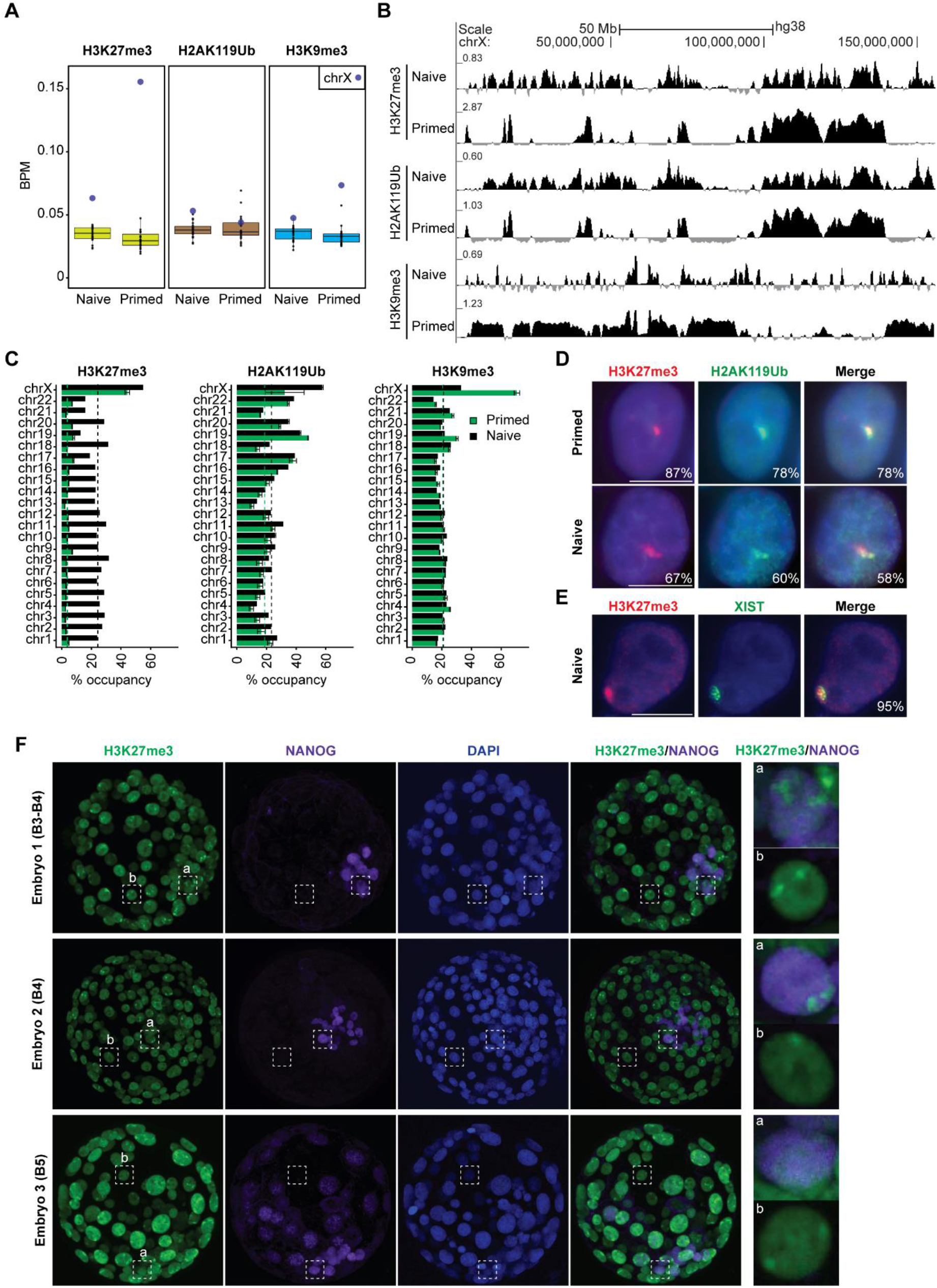
PRC-associated repressive histone modifications accumulate on dampened X chromosome in a XIST-dependent manner in naïve hESCs. **A)** Mean enrichment of H3K27me3, H2AK119Ub and H3K9me3 per chromosome in naïve and primed H9 hESCs. All individual data points are shown. **B)** CUT&RUN profiles (log2 enrichment over IgG) for H3K27me3, H2AK119Ub and H3K9me3 on the X chromosome in naïve and primed H9 hESCs. **C)** Percentage of H3K27me3, H2AK119Ub, and H3K9me3 peak occupancy per chromosome for primed and naïve H9 hESCs. Black and green dotted lines represent median genomic occupancy. Error bars correspond to the SD of replicates (n=2). **D)** Representative immunofluorescence images for H3K27me3 and H2AK119Ub in primed and naïve H9 hESCs. Percentages of cells displaying the representative pattern are indicated. Scale bar = 10 µm. **E)** Representative immunofluorescence images for H3K27me3 coupled with XIST RNA-FISH in naïve H9 hESCs. Percentages of cells displaying the representative pattern are indicated. Scale bar = 10 µm. **F)** Immunofluorescence images for H3K27me3 and NANOG in human pre-implantation blastocysts (B3 till B5 blastocysts).

Coverage profiles showed a distinct distribution of H3K27me3, H2AK119Ub, and H3K9me3 on the X chromosome in naïve compared to primed hESCs (Figure 3B). Notably, H3K27me3 and H2AK119Ub covered around 60% of the X chromosome in naïve hESCs and 40 to 50% in primed hESCs while most of the X chromosome (>60%) was covered by H3K9me3 in primed hESCs *vs* 33% in naïve hESCs (Figure 3C). H3K27me3 and H2AK119Ub patterns were highly correlated together and anticorrelated with H3K9me3 in both primed and naïve hESCs (Supplementary Figure 3A). H3K27me3 and H2AK119Ub signals accumulated into several smaller domains which spanned most of the XIST-coated dampened X chromosome in naïve hESCs (Figure 3B). We observed similar profiles of H3K27me3 and H2AK119Ub using other published datasets of naïve H9 hESCs reprogrammed using PXGL or T2iLGo (Supplementary Figure 3B, C). In contrast, H3K27me3 and H2AK119Ub accumulated over several large domains of the Xi, with the major one spanning around 50 Mb in the middle of the long arm, while H3K9me3 signal spanned most of the short arm as well as the most proximal and distal parts of the long arm (Figure 3B). Altogether, this showed that X chromosomes of naïve hESCs are enriched for repressive histone modifications, which are massively redistributed along the X upon transition from primed to naïve hESCs.

Patterns of H3K27me3 and H2AK119Ub in naïve hESCs were highly correlated to that of XIST in naïve as in primed hESCs, suggesting that the deposition of polycomb group associated histone modifications on the X chromosome is mediated by XIST in both contexts (Supplementary Figure 3D). We confirmed this hypothesis by showing a global loss of H3K27me3 and H2AK119Ub on the X chromosome in XIST KO and dox-treated CRISPRi clones compared to their WT counterpart (Supplementary Figure 3E), while H3K9me3 distribution and levels remained unchanged.

Immunofluorescence approaches confirmed CUT&RUN data (Figure 3D): one focus of H3K27me3 and H2AK119Ub was observed in > 60% of naïve H9 hESCs, with frequent co-occurrence of the 2 marks (Figure 3D). This H3K27me3 focal accumulation corresponds to the XIST-coated dampened X chromosome, as shown by immuno-RNA-FISH (Figure 3E). In the absence of XIST, H3K27me3 and H2AK119Ub co-accumulation was lost on the X chromosome in more than 90% of cells (Supplementary Figure 3F, G).

Since conflicting results existed regarding the accumulation of PRC-mediated repressive histone modifications on the X chromosomes prior to XCI establishment, we re-examined human pre-implantation embryos (Okamoto et al., 2011; Sahakyan et al., 2017; Vallot et al., 2017; van den Berg et al., 2009). Four blastocysts were marked with antibodies against NANOG and H3K27me3. In all embryos, we could observe cells with one or two H3K27me3 foci, both in NANOG+ and NANOG-cells (Figure 3F and Supplementary Figure 3H, I). The presence of two foci in a fraction of NANOG+ cells in 3 out of 4 embryos strongly suggest that active X chromosomes are indeed enriched for H3K27me3 in a fraction of epiblast cells of female blastocysts. Embryo 4, in which only one H3K27me3 focal enrichment is detected in around 10% of cells, is likely male (Supplementary Figure 3H, I). Overall, these data indicate that PRC-mediated repressive histone modifications are enriched on and broadly cover XIST-associated X chromosomes in most cells of female pre-implantation embryos and naïve hESCs.

### Highly expressed genes escape XIST-mediated dampening

The broad distribution of XIST and of repressive histone modifications along the dampened X chromosome led us to explore whether all X-linked genes are equally responsive to XIST-mediated regulation in naïve hESCs, based on their allelic expression profiles. Most of the X-linked genes that could be analyzed using SNPs are “XIST-sensitive” genes (78%), in agreement with XIST covering most of the X chromosome in naïve hESCs. These genes display lower expression from the XIST-associated X chromosome and become equally expressed from both X chromosomes in the absence of XIST (Figure 4A). In addition, we could identify a fraction of “XIST-resistant” genes (22%), with a pXi allelic ratio of ∼0.5 in the presence or absence of XIST, indicating equal expression from the dampened (pXi) and active (pXa) X chromosomes in naïve hESCs (Figure 4A).

**Figure 4:**
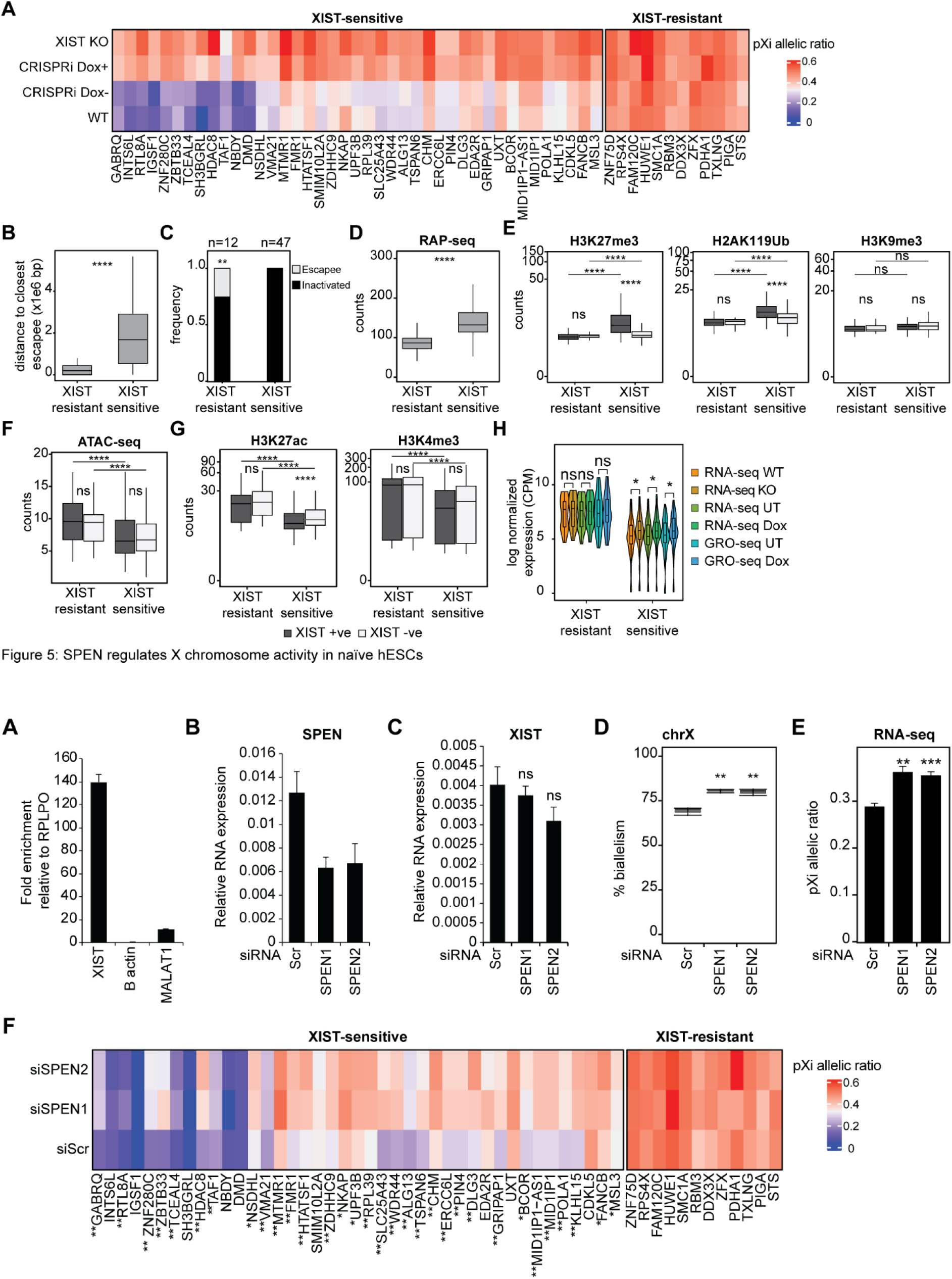
Highly expressed genes escape XIST-mediated dampening. **A)** Heat map showing the pXi allelic ratio of XIST-sensitive and XIST-resistant gene in WT, XIST KO, XIST CRISPRi UT (Dox-) and Dox-treated (Dox+) naïve H9 cells. **B)** Distance to closest escapee of XIST-sensitive and resistant genes. **C)** Frequency of escapees and inactivated X-linked genes as determined in primed H9 among XIST-sensitive and resistant genes. Chi-square test p-values: **<0.01. **D)** XIST RAP-seq counts on TSS±5kb of XIST-sensitive and resistant genes in naïve H9 cells. **E)** H3K27me3, H2AK119Ub, and H3K9me3 counts on TSS±5kb of XIST-sensitive and resistant genes in the presence (WT and untreated XIST CRISPRi) or absence (KO and Dox-treated XIST CRISPRi) of XIST in naïve H9 hESCs. **F)** ATAC-seq counts on TSS±5kb of XIST-sensitive and resistant genes in the presence (WT and untreated XIST CRISPRi) or absence (KO and Dox-treated XIST CRISPRi) of XIST in naïve H9 hESCs. **G)** H3K27ac and H3K4me3 counts on TSS±5kb of XIST-sensitive and resistant genes in the presence (WT and untreated XIST CRISPRi) or absence (KO and Dox-treated XIST CRISPRi) of XIST in naïve hESCs. **H)** Violin plots showing the log CPM expression level of XIST-sensitive and resistant genes in H9 naïve WT, KO, XIST CRISPRi UT and Dox-treated cells, obtained from RNA-seq and fastGRO-seq data. Unless otherwise specified, p-values were calculated by two-sided unpaired t-test. Levels of significance: ns≥0.05; * < 0.05; ** < 0.01; *** < 0.001; **** < 0.0001.

While no significant difference was observed in the distance to the XIST locus between XIST-resistant and XIST-sensitive genes (Supplementary Figure 4A), XIST-resistant genes tended to be located in the vicinity of XCI escapees and were enriched on the short arm, while XIST-sensitive genes appeared to be scattered along the X (Figure 4B and Supplementary Figure 4C). Of note, all the genes that escape XCI in primed hESCs appeared to be XIST-resistant in naïve hESCs (Figure 4C), suggesting that this set of genes is refractory to XIST repressive activities (dampening or inactivation), whatever the context.

XIST-sensitive genes were significantly more contacted by XIST RNA and had higher accumulation of H3K27me3 and H2AK119Ub compared to XIST-resistant genes, while H3K9me3 and DNA methylation levels were similar in both categories (Figure 4D, E and Supplementary Figure 4C). Accordingly, only XIST-sensitive genes display reduced H3K27me3 and H2AK119Ub levels in the absence of XIST, with levels reaching that of XIST-resistant genes, while H3K9me3 remained constant across gene categories and genotypes (Figure 4E and Supplementary Figure 4D).

Conversely, the TSS of XIST-sensitive genes were less accessible than that of XIST-resistant genes and were marked by lower levels of active histone modifications H3K27ac and H3K4me3 (Figure 4F, G). H3K27ac, but not H3K4me3 levels were elevated specifically at XIST-sensitive genes upon loss of XIST but remained nevertheless lower than XIST-resistant genes. This indicates that, while XIST is responsible for the deposition of repressive histone marks at target genes, it only mildly prevents the acquisition of active promoters’ features. In agreement with these observations, XIST-sensitive genes were expressed at lower levels than XIST-resistant genes in naïve hESCs and do not reach expression levels of XIST-resistant genes in the absence of XIST (Figure 4H). Interestingly, a similar tendency was observed in pre- and post-implantation human embryos (Supplementary Figure 4E). Altogether, these results show that, in naïve hESCs, and possibly in pre-implantation embryos, XIST attenuates the expression of the majority of X-linked genes, with highly expressed genes resisting XIST targeting and dampening activity.

### SPEN is involved in XIST-mediated dampening

Previous studies have shown that SPEN interacts with the A repeat of mouse Xist RNA and is rapidly recruited to the X chromosome at the initiation of XCI to promote histone deacetylation (Chu et al., 2015; Dossin et al., 2020; McHugh et al., 2015). Nevertheless, whether SPEN interacts with human XIST and is recruited on the dampened X chromosome in naïve hESCs is unknown. We first analyzed SPEN expression and protein abundance by RNA-seq and mass spectrometry, respectively. SPEN mRNA levels were 2.5-fold higher in naïve compared to primed H9 hESCs (Supplementary Figure 5A). However, SPEN protein abundance appears largely similar between naïve and primed H9 hESCs (Supplementary Figure 5B). RNA immunoprecipitation (RIP) experiments using SPEN antibody led to a 140-fold enrichment for XIST RNA relative to control RPLPO RNA. In contrast, β-actin mRNAs or MALAT1 non-coding RNAs that were not reported to interact with SPEN were not significantly immuno-precipitated (Figure 5A). This strongly suggests that SPEN preferentially associates with XIST in naïve hESCs.

**Figure 5:** SPEN is involved in XIST-mediated dampening. **A)** RTqPCR showing fold enrichment levels of XIST, β-actin, and MALAT1 normalized to RPLPO following RNA immunoprecipitation (RIP) of SPEN (n=3). Error bars correspond to the SD of replicates. **B)** RTqPCR analysis of SPEN expression after siRNA KD using two different mixes of siRNAs SPEN1 and SPEN2 (n = 3). Values are normalized to β-actin. Error bars correspond to the SD of replicates. **C)** RTqPCR analysis of XIST expression after SPEN KD (n = 3). Values are normalized to β-actin. Error bars correspond to the SD of replicates. Wilcoxon p-value = ns≥0.05. **D)** Percentage of biallelically expressed SNPs from X chromosome in scramble and SPEN KD naïve H9 cells. **E)** pXi allelic ratio from RNA-seq data of scramble and SPEN KD naïve H9 hESCs (n=3). Error bars correspond to the SD of replicates. **F)** Heatmap showing the pXi allelic ratio of XIST-sensitive and resistant gene expression upon SPEN KD in naïve H9 hESCs. An increase in pXi allelic ratio (FC>1.2) from one or both siSPEN mixes is indicated by * or ** respectively. Unless otherwise specified, p-values were calculated by two-sided unpaired t-test. Levels of significance: ns≥0.05; * < 0.05; ** < 0.01; *** < 0.001; **** < 0.0001.

We then transiently knocked down SPEN in naïve H9 hESCs using two different mixes of siRNAs (SPEN1 and SPEN2) (Figure 5B). SPEN knockdown had no significant impact on pluripotency factor or XIST expression level (Supplementary Figure 5C and Figure 5C). Differential expression analysis revealed that 775 genes (351 upregulated and 424 downregulated; log2FC > |1| and FDR<0.05) or 652 genes (379 upregulated and 273 downregulated; log2FC > |1| and FDR<0.05) showed altered expression levels upon siSPEN1 or siSPEN2 KD compared to control siScr (Supplementary Figure 5D, E). DEGs were distributed on all chromosomes and not specifically enriched on the X chromosome, consistent with the role of SPEN as a regulator of different key pathways (Giaimo et al., 2021; Kuroda et al., 2003; Shi et al., 2001) (Supplementary Figure 5F).

To evaluate the role of SPEN in the regulation of X chromosome activity, we analyzed the percentage of biallelically detected X-linked SNPs in SPEN KD naïve hESCs compared to control cells by RNA-seq. We observed a slight increase of biallelically expressed SNPs upon SPEN KD, suggesting that SPEN loss leads to higher expression of a subset of X-linked genes, presumably from the XIST-coated X chromosome (Figure 5D and Supplementary Figure 5G). This was confirmed by calculating the pXi allelic ratio, which was significantly increased in the SPEN KD condition compared to siScr (Figure 5E), albeit to a lesser extent than that in XIST null conditions (Figure 1J). This can be due to incomplete SPEN KD or to the involvement of other factors in dampening, and likely explains why the X/A ratio was not affected upon SPEN KD (Supplementary Figure 5H).

Finally, we explored the dependency of XIST-sensitive and XIST-resistant genes on SPEN activity. Interestingly, 32 out of 41 XIST-sensitive genes (78%) displayed higher pXi allelic ratio in at least one or both siSPEN compared to siScr control (Figure 5F). In contrast, no change in pXi allelic ratio was observed for XIST-resistant genes in the absence of SPEN, which is consistent with XIST recruiting SPEN to X-linked genes to exert its repressive activity (Figure 5F). Overall, these results reveal the implication of SPEN in mediating the dampening activity of XIST in naïve hESCs.

## Discussion

The intriguing uncoupling between XIST accumulation and XCI during early human development raises several fundamental questions that we aimed to address: Are there mechanisms protecting the X chromosome from being silenced? is XIST functional? is XIST involved in alternative dosage compensation mechanisms? If so, what are the underlying mechanisms?

We had previously hypothesized that the lncRNA XACT could be an antagonist of XIST during pre-implantation development, when it co-accumulates with XIST on active X chromosomes (Vallot et al., 2017). In that case, loss of XACT expression in naïve hESCs, which mimics cells of the pre-implantation epiblast, should lead to inactivation of XIST-coated X chromosomes. As only one active X is decorated by XIST in these cells (the previous Xi), XIST-mediated inactivation should be compatible with cell viability, allowing us to test our hypothesis. By deleting a large genomic fragment encompassing all XACT transcription start sites, we show that XACT does not control X chromosome activity in naïve hESCs. In addition, few genes were differentially expressed in XACT KO clones, indicating that XACT is not a potent regulator of gene expression in naïve hESCs. This, however, does not exclude that XACT might be functional in other cellular contexts. Previous study showed that insertion of a XACT transgene in mouse cells influences XIST accumulation *in cis* (Vallot et al., 2017), which suggests a potential function of XACT during the initiation phase of XCI by dictating the choice of the X to be inactivated. Moreover, XACT deletion in primed hESCs affects neural differentiation, through mechanisms that remain to be determined (Motosugi et al., 2021). Finally, XACT is highly expressed in primordial germ cells (Chitiashvili et al., 2020). This could reflect either a cellular context resembling naïve pluripotency, permissive for XACT expression, or a more specific role for XACT during germ line specification.

An alternative scenario to explain why XACT KO does not result in XCI in naïve hESCs invokes the production of a non-functional form of XIST at this stage. We could discard this hypothesis by demonstrating that XIST triggers deposition of polycomb-mediated histone modifications, interacts with SPEN and modulates the activity of the coated X chromosome. Indeed, we reveal an enrichment in H3K27me3 and H2AK119ub on active X chromosomes in naïve hESCs and in human pre-implantation blastocysts. This contrasts with previous report where active X chromosomes were found, by IF, devoid of H3K27me3 in human embryos (Okamoto et al., 2011). Such discrepancy may result from different embryo handling and/or hybridization conditions. Alternatively, enrichment of active Xs in polycomb-mediated histone modifications may be transient or heterogeneous, and thus variably captured, in agreement with foci of H3K27me3 being visible only in a subset of blastocyst nuclei within a given embryo. These observations are nevertheless in agreement with recent findings in cynomolgus monkey pre-implantation embryos, in which H3K27me3 marks both active X chromosomes (Okamoto et al., 2021). In the latter study, Okamoto and colleagues also unraveled the dynamics of XCI, with earlier establishment in the trophectoderm compared to embryonic lineages (Okamoto et al., 2021). Interestingly, we observe a nonnegligible number of NANOG-cells, which correspond to trophectoderm cells, harboring one focus of H3K27me3, which could indicate that XCI initiates prior to implantation in human, as in cynomolgus monkey.

Even if active and inactive X chromosomes feature enrichment in polycomb-repressive marks, our CUT&RUN data reveal a redistribution of the X chromosome repressive chromatin landscape upon transition from primed to naïve hESCs. Since we are comparing active and inactive X chromosome features in an isogenic context, we can confidently link the observed differences to the cellular state. Repressive modifications were broadly distributed on active X chromosome in naïve cells, but their enrichment was higher on the inactive X of primed hESCs. We demonstrated that XIST is responsible for the deposition of these modifications in naive hESCs, on the basis of the following observations: (i) only the XIST-coated chromosome displays such enrichment; (ii) H3K27me3 and H2AK119Ub patterns on active X follow that of XIST, as probed by RAP-seq and (iii) loss of XIST (through KO or inducible CRISPRi) leads to loss of H3K27me3 and H2AK119Ub specifically on the X chromosome.

The redistribution of XIST on active X chromosome upon hESC resetting, which leads to a large proportion of the naïve active X chromosome being contacted by XIST, is consistent with previous findings of XIST clouds being more dispersed in naïve hESCs and human pre-implantation embryos, as observed in RNA-FISH experiments (Sahakyan et al., 2017; Vallot et al., 2017). How XIST propagates on the X chromosome and what drives such distinct XIST distribution in hESCs is still unknown. It has been proposed that transposable elements, and in particular LINEs, might serve as waystations for XIST propagation on the X chromosome (Bailey et al., 2000; Cantrell et al., 2009). Here we observed a slight correlation between XIST coverage and the density of LINE and retrotransposons in naïve hESCs. Thus, upon zygotic genome activation, the newly produced XIST RNA molecules might be preferentially targeted to TE-enriched regions. Spatial organization of the X chromosome may also orchestrate XIST distribution. This hypothesis is based on previous studies performed in the mouse, whereby Xist exploits the three-dimensional genome architecture to spread across the X chromosome during XCI initiation (Engreitz et al., 2013).

XIST distribution has consequences for X chromosome activity. Indeed, we show that XIST is responsible for attenuating expression level of genes that its RNA contacts. Thus, broad distribution of XIST results in most X-linked genes being dampened. There are, however, genes that are less contacted by XIST and whose expression is not attenuated. We show that genes escaping dampening are expressed at higher levels compared to XIST-sensitive genes, not only in naïve hESCs, but also in embryos. It has been previously reported that genes escaping XCI are also expressed at higher levels than other X-linked genes, which was proposed to be connected to the strong purifying selection characterizing escapees (Slavney et al., 2016). The fact that dampening resistant genes are also escaping XCI or located in proximity to XCI escapees suggest common mechanisms for resisting XIST repressive activities. Reversely, our data, which reveals that dampening is mediated at the level of transcription initiation, involves the XIST interactor SPEN and is associated with XIST-mediated deposition of PRC modifications, suggest a conserved mode of action for XIST in both contexts.

The question remains as to what is preventing full silencing of X chromosomes at this stage. Multiple non-exclusive scenarios could be envisioned, such as the absence and/or defects (in expression or post-translational modifications) of certain XIST effectors. Another possible explanation is the presence or absence of post-transcriptional modifications on XIST RNA that can alter its interaction with protein partners or disrupt its folding. Finally, other unknown factors might prevent XIST from silencing the X chromosome. Whatever the mechanisms, signals that trigger the switch in XIST activity must be linked to developmental progression. This is reminiscent to mouse ESCs, where inducible XIST induces XCI but not to its completion, with differentiation being required for full XCI in a manner that involves SMCHD1 (Bowness et al., 2022).

Finally, one might question the functional relevance of dampening. We could not observe any change in cell morphology, proliferation and transcriptomic signature when XIST was deleted or repressed in naïve hESCs, suggesting that X chromosome dampening is not required for stem cell fitness *in vitro*. Whether it is necessary for progression of peri-implantation development, and whether altered dampening could leave scars and affect future development is unknown. “Pre-marking” the X chromosome with polycomb repressive modifications and attenuating X chromosome activity could facilitate initiation of XCI, providing asymmetry between the 2 Xs is subsequently installed. It is yet unclear whether dampening is a hominoid-specific mechanism or whether it is deployed in all primates. Recent investigation in cynomolgus monkey pre-implantation embryos revealed a transient dampened-like status, which however did not lead to complete X chromosome dosage compensation, as X-linked gene expression levels remained higher in females compared to males (Okamoto et al., 2021). Of note, X chromosome dampening is the dosage compensation strategy at stake in *Caenorhabditis elegans* where it is achieved by the dosage compensation complex (DCC) (Strome et al., 2014). Overall, this highlight both diversity and convergence in the evolution of mechanisms and actors underlying X chromosome dosage compensation across species and clades, with dampening being triggered by unrelated sets of factors in nematode and mammals, while a XIST ribonucleoprotein complex displays distinct activities according to the cellular context in a given species.

## Acknowledgments

We thank lab members for critical evaluation of the work leading to this publication, and Céline Morey and Slimane Ait Si Ali for critical reading of the manuscript. We thank the Epigenomic, the Microscopy, the Vectorology and the BIBS Platforms, all hosted in UMR7216 Epigenetic and Cell Fate, for technical advice and access to instruments. We thank the ICGex NGS platform of the Institut Curie supported by the grants ANR-10-EQPX-03 (Equipex) and ANR-10-INBS-09-08 (France Génomique Consortium) from the Agence Nationale de la Recherche (“Investissements d’Avenir” program), by the ITMO-Cancer Aviesan (Plan Cancer III) and by the SiRIC-Curie program (SiRIC Grant INCa-DGOS-465 and INCa-DGOSInserm_12554) for the high-throughput sequencing. We thank the Bioinformatics platform of the Institut Curie for data management, quality control and primary analysis. This project was funded by Agence Nationale pour la Recherche [ANR-14-CE10-0017 to C.R.]; European Research Council (ERC) under the European Union’s Horizon 2020 research and innovation program [101020423]; LabEx ‘Who Am I?’ [ANR-11-LABX-0071]; Université de Paris IdEx [ANR-18-IDEX-0001] funded by the French Government through its ‘Investments for the Future’ program. E.C. was supported by fellowships from the French Ministry of Education and Research and from the French Medical Research Foundation (FRM).

## Ethics declarations

The authors declare no competing interests.

## Methods

### Human cell lines and culture conditions

Experiments were carried out using the female H9 hESCs obtained from the WiCell Research Institute. Research on human embryonic stem cells has been approved by Agence de la Biomédecine and informed consent was obtained from all subjects.

Primed H9 hESCs were cultured on Matrigel-coated culture dishes in mTeSR™1 media (Stemcell Technologies) according to the manufacturer instructions, in 5% O2 and 5% CO2 at 37°C. They were routinely passaged in clumps using gentle cell dissociation reagent (Stemcell Technologies) according to the manufacturer instructions. For experiments requiring single-cell suspension, cells were incubated with Accutase (Stemcell Technologies) and plated in fresh mTeSR™1 media supplemented with 10 μM of Y-27632 (Stemcell Technologies).

Naïve H9 hESCs were generated by chemical resetting of the H9 primed hESCs using NaiveCult (Stemcell Technologies) or PXGL protocol, as previously described, and cultured on inactivated mouse embryonic fibroblasts (MEFs) (Bredenkamp et al., 2019; Guo et al., 2017). PXGL naïve hESCs were cultured in PXGL medium consisting of a 1:1 mixture of DMEM/F12 (Sigma-Aldrich) and Neurobasal media supplemented with 0.5% N2 supplement, 1% B27 supplement, 2mM l-glutamine, 100 µM β-mercaptoethanol and 1x penicillin– streptomycin (all from Gibco, Thermo Fisher Scientific) as well as 1μM PD0325901 (Axon Medchem, 1408; CAS: 391210-10-9), 2μM XAV939 (Cell guidance systems, SM38-10; CAS: 284028-89-3), 2μM Gö6983 (Tocris, 2285; CAS: 133053-19-7), 10 ng/mL human LIF (Peprotech, 300-05). Naïve hESCs were routinely passaged as single cells every 3 days at a ratio of 1:3 using TrypLE™ Express (1X) (Gibco, Thermo Fisher Scientific) and plated in fresh media supplemented with 10 μM of Y-27632 (Stemcell Technologies).

### Human pre-implantation embryos

The use of human embryo donated to research as surplus of IVF treatment was allowed by the French embryo research oversight committee: Agence de la Biomédecine, under approval number RE18-010R. All human pre-implantation embryos used in this study were obtained from and cultured at the Assisted Reproductive Technology unit of the University Hospital of Nantes, France, which are authorized to collect embryos for research under approval number AG110126AMP of the Agence de la Biomédecine. Embryos used were initially created in the context of an assisted reproductive cycle with a clear reproductive aim and then voluntarily donated for research once the patients have fulfilled their reproductive needs or tested positive for the presence of monogenic diseases. Informed written consent was obtained from both parents of all couples that donated spare embryos following IVF treatment. Before giving consent, people donating embryos were provided with all of the necessary information about the research project and opportunity to receive counselling. No financial inducements are offered for donation. Molecular analysis of the embryos was performed in compliance with the embryo research oversight committee and The International Society for Stem Cell Research (ISSCR) guidelines (Kimmelman et al., 2016).

Human embryos were thawed following the manufacturer’s instructions (Cook Medical: Sydney IVF Thawing kit for slow freezing and Vitrolife: RapidWarmCleave or RapidWarmBlast for vitrification). Human embryos frozen at 8-cell stage were loaded in a 12-well dish (Vitrolife: Embryoslide) with non-sequential culture media (Vitrolife G-TL™) under mineral oil (Origio: Liquid Paraffin), at 37°C, in 5% O2 and 6% CO2.

### Generation of XACT and XIST KO cell line

Single guide RNA (sgRNA) sequences flanking XACT and XIST promoters and first exon were obtained using the web-based tool CRISPOR (http://crispor.tefor.net/) and are provided in the Supplementary Table 3. sgRNAs were cloned under an U6 promoter into the pSpCas9(BB)−2A-GFP (a gift from Feng Zhang, Addgene #48138) and the pSpCas9(BB)-2A-mCherry (generated in house, by replacing the green fluorescent protein (GFP) with a mCherry reporter using the NEBuilding HiFi DNA Assembly Cloning Kit (New England Biolabs) (Ran et al., 2013). Using the Amaxa 4D-NucleofectorTM system (Lonza), 1 million H9 primed hESCs were transfected with 2.5 µg of each plasmid (to a total of 5 µg). Cells were sorted by fluorescence-activated cell sorting (INFLUX 500-BD BioSciences) 48 h after transfection. Double-positive cells were plated into a matrigel-coated 6 cm petri dish in mTeSR supplemented with 1x CloneR (Stemcell Technologies). Individual colonies were manually picked into 96 wells ∼10 days after transfection. Deletions and inversion events were screened by PCR. Primer sequences can be found in Supplementary Table 4. XACT KO and XIST KO H9 primed hESCs were reprogrammed to naïve using NaiveCult and PXGL protocols, respectively, as described above.

### CRISPR inhibition

sgRNA targeting XIST promoter was designed using the web-based tool CRISPOR (http://crispor.tefor.net/) and cloned into the PB_rtTA_BsmBI vector (a gift from Mauro Calabrese, Addgene, #126028) (Schertzer et al., 2019). sgRNA sequence can be found in Supplementary Table 3. Using the Amaxa 4D-NucleofectorTM system (Lonza), 1 million H9 primed hESCs were transfected with 1.5 µg of PB_tre_dCas9_KRAB (a gift from Mauro Calabrese, Addgene, #126030), 0.75 µg of PB_rtTA_BsmBI and 1.5 µg of piggyBac transposase (Wang et al., 2008). Cells were then treated with G418 (350 µg/mL) and hygromycin (350 µg/mL) until separate colonies were obtained. The number of random insertions in the genome was verified by qPCR and clones with the lowest insertion number (n=2) were used for further experiments. For XIST depletion, PXGL-reprogrammed CRIPSRi naïve hESCs were treated with 1µg/mL doxycycline for 10 days.

### CUT&RUN

CUT&RUN was performed as previously described (Skene and Henikoff, 2017). Briefly, 0.3 million cells per replicate were bound to 20 uL concanavalin A-coated beads (Bangs Laboratories) in binding buffer (20 mM HEPES, 10 mM KCl, 1 mM CaCl2 and 1 mM MnCl2). The beads were washed and resuspended in Dig-wash buffer (20 mM HEPES, 150 mM NaCl, 0.5 mM spermidine, 0.05% digitonin). The primary antibodies (1:50) were added to the bead slurry and rotated at RT for 1 hour. The beads were washed by Dig-wash buffer and pA-Mnase fusion protein (1:400, produced by the Institut Curie Recombinant Protein Platform, 0.785mg/mL) was added and rotated at RT for 15 min. After two washes, the beads were resuspended in 150 µL Dig-wash buffer, and the MNase was activated with 2 mM CaCl2 and incubated for 30 min at 0 °C. MNase activity was terminated with 150 µL 2XSTOP (200 mM NaCl, 20 mM EDTA, 4 mM EGTA, 50 µg/ml RNase A and 40 µg/ml glycogen). Cleaved DNA fragments were released by incubating for 20 min at 37 °C, followed by centrifugation for 5 min at 16,000g at 4 °C and collection of the supernatant from the beads on a magnetic rack. The DNA was purified by phenol:chloroform and libraries were prepared using the TruSeq ChIP Library Preparation Kit from Illumina following the manufacturer’s protocol, and sequenced on a NovaSeq 6000 instrument (ICGex - NGS platform, Paris, France) to generate 2X100 paired-end reads. All the antibodies used in this study are listed in the Supplementary Table 5.

### RAP-seq

RNA antisense purification followed by DNA sequencing (RAP-seq) for human XIST was performed as previously described (Engreitz et al., 2013). The human XIST oligo pool was ordered from GenScript and amplified as previously described to generate ssDNA biotinylated oligos (Engreitz et al., 2015). The oligo sequences are listed in the Supplementary Table 6. Briefly, 20 million cells were harvested and incubated with 10 mL of freshly-made 2mM DSG at RT for 45 min. Cells were further crosslinked with 10 mL 3% formaldehyde for 10 min at 37°C, and the reaction was stopped by adding glycine to a final concentration of 500 mM. Cells were pelleted at 4°C and stored at 80°C. Nuclei were isolated in cold lysis buffer (20 mM HEPES; pH 7.5, 50 mM KCl, 1.5 mM MnCl2, 1 % IGEPAL CA630 (NP-40), 0.4 % sodium deoxycholate, and 0.1 % N -lauroylsarcosine). Chromatin was solubilized by sonication and fragmented by TURBO DNase digestion (Thermo Fisher Scientific). XIST pulldown was performed on 10 million cells using 1 µg of Streptavidin Dynabeads™ MyOne™ C1 (Thermo Fisher Scientific) and 50 pmol of biotinylated oligos. DNA was eluted by RNase H digestion, and crosslinking was reversed via proteinase K digestion at 65°C for 1h. Finally, DNA was purified using the GeneJET Gel Extraction kit (Thermo Fisher Scientific) and libraries were prepared using the TruSeq ChIP Library Preparation Kit from Illumina, and sequenced on a NovaSeq 6000 instrument (ICGex - NGS platform, Paris, France) to generate 2X100 paired-end reads.

### ATAC-seq

ATAC-seq was performed as previously described (Buenrostro et al., 2015). Briefly, 50.000 cells were resuspended in 50µL cold lysis buffer (10mM Tris-HCl; pH 7.4, 10mM NaCl, 3 mM MgCl2, 0.1 % Igepal CA-630) and centrifuged for 10 min at 500g at 4°C. The nuclei pellets were resuspended in 50 μl transposase reaction mix (25 µl 2X TD buffer, 2.5 µl transposase, and 22.5 µl H2O) and incubated at 37°C for 30 min in a thermomixer with 1000 RPM mixing. Reactions were cleaned up using the MinElute PCR purification kit (QIAGEN) and DNA was eluted in 10 µ elution buffer. Transposed DNA were pre-amplified for 5 cycles in 50 µl reaction mix (2.5 μl of 25 μM primer Ad1, 2.5 μl of 25 μM primer Ad2, 25 μl of 2X Master Mix, 10 µL H2O and 10 μl transposed elution) at the following cycling conditions: 72°C, 5 min; 98°C, 30 s; then 5 cycles of (98°C, 10 s; 63°C, 30 s; 72°C, 1 min). Then, 15 μl of qPCR amplification reaction (5 μl of pre-amplified sample; 0.5 μl of 25 μM primer Ad1, 0.5 μl of 25 μM primer Ad2, 5 μl of 2X NEBNext Master Mix, 0.24 μl of 25x SYBR Green in DMSO, and 3.76 μl of H2O) was carried out at the following cycling conditions: 98°C, 30 s; then 20 cycles of (98°C, 10 s; 63°C, 30 s; 72°C, 1 min). The required number of additional cycles for each sample was calculated. After the final amplification, double-sided bead purification was performed with AMPure XP beads. Final ATAC-seq libraries were eluted in 20 μl nuclease-free H2O from the beads and were sequenced on a NovaSeq 6000 instrument (Novogene, UK) to generate 2X150 paired-end reads.

### RNA-seq

Total RNAs were collected using RNeasy mini kit (QIAGEN) and extracted following the manufacturer’s recommendation. Libraries were generated using the Illumina Stranded Total RNA Prep Ligation with Ribo-Zero Plus kit according to the manufacturer’s recommendation and sequenced on a NovaSeq 6000 instrument (ICGex - NGS platform, Paris, France) to generate 2X100 paired-end reads.

### fastGRO-seq

Low input fastGRO-seq was performed as previously described (Barbieri et al., 2020). Nuclei were isolated from 5 million cells and nuclear run-on was performed by adding 25 µL of 2X Nuclear run-on buffer (10 mM Tris-HCl; pH 8, 5 mM MgCl2, 300 mM KCl, 1 mM DTT, 500 μM ATP, 500 μM GTP, 500 μM 4-thio-UTP, 2 μM CTP, 200 μ/ml Superase-in, 1% Sarkosyl (N-Laurylsarcosine sodium salt solution) at 30°C for 7 min. RNA was extracted with TRIzol LS reagent (Invitrogen) and quantified using the nanodrop 2000. 30 µg of RNA were fragmented by sonication and the efficiency was analyzed using the Agilent High Sensitivity RNA ScreenTape Assay. Fragmented RNA was incubated in Biotinylation Solution (25 mM HEPES; pH 7.4, 10 mM EDTA; pH 8.0, 50 µg MTS-Biotin (Biotium)) for 30 min in the dark at 24°C and 800 rpm. RNA was then precipitated using ethanol and resuspended in nuclease-free water. After DNase treatment, biotinylated RNA was enriched using M280 Streptavidin Dynabeads (Invitrogen) and precipitated using ethanol. Libraries were prepared using the Illumina Stranded Total RNA Prep Ligation with Ribo-Zero Plus kit according to the manufacturer’s recommendation and sequenced on a NovaSeq 6000 instrument (ICGex - NGS platform, Paris, France) to generate 2X100 paired-end reads.

### MeD-seq

MeD-seq assays were essentially performed as previously described (Boers et al., 2018). Briefly, 10 µl genomic DNA (input 90 ng) from naïve hESCs were digested with LpnPI (New England Biolabs) generating 32 bp fragments around the fully methylated recognition site containing a CpG. These short DNA fragments were further processed using the ThruPlex DNA–seq 96D kit (Rubicon Genomics Ann Arbor). Stem-loop adapters were blunt-end ligated to repaired input DNA and amplified to include dual indexed barcodes using a high-fidelity polymerase to generate an indexed Illumina NGS library. The amplified end product was purified on a Pippin HT system with 3% agarose gel cassettes (Sage Science). Multiplexed samples were sequenced on Illumina NextSeq2000 systems for paired-end read of 50 bp according to the manufacturer’s instructions. Dual indexed samples were de-multiplexed using bcl2fastq software (Illumina).

### RNA-FISH

Cells preparation: Primed hESCs were grown on coverslips. Naïve hESCs were centrifuged onto Superfrost Plus slides (VWR) using the Cytospin 3 Cytocentrifuge (Shandon). The cells were fixed for 10 min in a 3% paraformaldehyde solution (Electron Microscopy Science) and permeabilized for 5–10 min in ice-cold CSK buffer (10 mM PIPES; 300 mM sucrose; 100 mM NaCl; 3 mM MgCl2; pH 6.8) supplemented with 0.5% Triton X-100 (Sigma-Aldrich) and 2 mM VRC (New England Biolabs).

Probes preparation: RNA-FISH probes were obtained after Nick translation of fosmids/BAC constructs purified using phenol:chloroform: 1 μg of purified DNA was labelled for 3 h at 15°C with fluorescent dUTPs (SpectrumOrange and SpectrumGreen from Abott Molecular and Cy5-UTPs from GE HealthCare Life Science).The templates used in this study are: human XIST fosmid (Bacpac resource center, WI2-3059D20), human POLA1 BAC (Bacpac resource center, RP11-11104L9), human XACT BAC (Bacpac resource center,RP11135D3) and human HUWE1 BAC (Bacpac resource center, RP11-975N19).

Hybridization: 100 ng of probes were supplemented with 1μg of Cot-I DNA (Invitrogen) and 3μg of Sheared Salmon Sperm DNA (Invitrogen). After precipitation, the probes were resuspended in deionized formamide (Sigma Aldrich), denatured for 7 min at 75°C and further incubated for 10 min at 37°C. Probes were mixed with an equal volume of 2× Hybridization Buffer (4× SSC, 20% dextran sulfate, 2 mg/ml BSA, 2 mM VRC). Coverslips were dehydrated in 80–100% ethanol washes and incubated with the hybridization mix at 37°C overnight in a humid chamber. Next, the coverslips were washed for 4 min at 42°C three times with 50% formaldehyde/2× SSC (pH 7.2) and three times with 2× SSC. The coverslips were mounted in Vectashield plus DAPI (Vector Laboratories).

### Immunofluorescence staining

Primed and naïve hESCs were prepared as described in the above section and fixed for 10 min in a 3% paraformaldehyde solution (Electron Microscopy Science). Cells were then permeabilized for 7 min with ice-cold PBS supplemented with 0.5% Triton X-100 (Sigma-Aldrich). Cells were blocked in 1X PBS/1% BSA (Sigma-Aldrich) for 15 min at RT and incubated for 45 min at RT with primary antibody diluted in 1X PBS/1% BSA. After 3 PBS washes, cells were incubated for 40 min at RT with Alexa Fluor 488 nm anti-rabbit or Alexa Fluor 594 nm anti-mouse secondary antibodies (Life Technologies). Finally, coverslips were mounted in Vectashield plus DAPI (Vector Laboratories). For combined Immunofluorescence and RNA-FISH, immunofluorescence staining was first performed then RNA-FISH as described above.

Human embryos were fixed at the B3, B4, or B5 stages according to the grading system proposed by Gardner and Schoolcraft (Gardner and Schoolcraft, 1999). Embryos were fixed with 4% paraformaldehyde (Electron Microscopy Sciences) for 5 min at RT and washed in 1X PBS/0,1% BSA. Embryos were permeabilized and blocked in 1X PBS/0.2% Triton/10% FBS at RT for 60 min. Samples were incubated overnight at 4°C with primary antibodies. After 3 PBS washes, embryos were incubated for 2h at RT with Alexa Fluor 488 nm anti-mouse or Alexa Fluor 568 nm anti-mouse and Alexa Fluor 647 nm anti-goat secondary antibodies (Life Technologies) along with DAPI counterstaining (Invitrogen). All primary antibodies used in this study are listed in the Supplementary Table 5.

### Microscopy and image analysis

Fluorescent microscopy images for hESCs were taken on a fluorescence DMI-6000 inverted microscope with a motorized stage (Leica), equipped with a CCD Camera HQ2 (Roper Scientifics) and an HCX PL APO 64-100X oil objective (numerical aperture, 1.4, Leica) using the Metamorph software (version 7.04, Roper Scientifics). Approximately 40 optical z-sections were collected at 0.3 μm steps, at different wavelengths depending on the signal (DAPI [360 nm, 470 nm], FITC [470 nm, 525 nm], Cy3 [550 nm, 570 nm], and Cy5 [647 nm, 668 nm]). We computed the dispersion of XIST RNA by comparing the cumulative volume of the signal to the theoretical spherical volume it could occupy based on the maximal radial distance. Embryo confocal immunofluorescence images were acquired with A1-SIM Nikon® confocal microscope and 20X oil objective. Whole embryos were imaged with 0,5-1 μm optical sections. Stacks were processed using ImageJ, and are represented as a 2D ‘maximum projection’ throughout the manuscript.

### Total RNA extraction and RTqPCR

Total RNAs were collected using RNeasy mini kit (QIAGEN) and extracted following the manufacturer’s recommendation. Quantification of the extracted RNAs was done using the nanodrop 2000. DNase step was performed on 1 μg of RNA for 30 min at 37°C using TURBO™ DNase (Thermo Fisher Scientific). RNAs were reverse transcribed using the Superscript IV kit (Thermo Fisher Scientific) following the manufacturer’s recommendation. cDNAs were diluted 2.5 times in water and RNA expression level was assessed by real time quantitative PCR (RTqPCR) using the Power SYBR Green Master Mix (Thermo Fisher Scientific) and ViiA-7 Real-Time PCR system (Applied Biosystems). Transcript RNA levels were normalized against β-actin reference gene following the 2-ΔCt method. All RT-qPCR primers used in this study are listed in the Supplementary Table 7.

### RNA immunoprecipitation

Cellular extract from 1 million H9 naïve hESCs was prepared by adding 1 mL HNTG buffer (20 mM Hepes; pH 7.9, 50 mM NaCl, 1% triton X-100, 1 mM MgCl2, 1 mM EDTA, 10% glycerol) followed by incubation on ice for 25 min. Cells were then centrifuged at 16,000 g 4°C for 25 min. The supernatant was collected and 80 µL was used as input fraction. 30 μL Magna ChIP™ protein G magnetic beads (Merck) per IP were washed with PBS-Tween 0.1% three times. Then 2.5 μg of SPEN antibody (Abcam, AB72266) were added to the beads followed by 10 min incubation at RT with rotation. Beads were washed three times with PBS-Tween 0.1% and resuspended with HNTG buffer. 30 μL of beads were added to the lysate prepared above and incubated for 1.5 hours at 4°C with rotation. After immunoprecipitation, beads were washed three time with 500 μL of HNTG buffer. For RNA preparation, Input and IP samples were treated with 200 μg of proteinase K in a total volume of 200 μL for 45 min at 65°C. RNA was purified using the RNeasy MinElute Cleanup kit (Qiagen) according to the manufacturer’s recommendations in a final volume of 14 μL of water. DNase step and reverse transcription were performed as described above. RNA enrichment level was assessed by RTqPCR using the Power SYBR Green Master Mix (Thermo Fisher Scientific) and ViiA-7 Real-Time PCR system (Applied Biosystems).

### siRNA knockdown SPEN

SPEN knockdown was achieved using two different mixes of siRNAs. Mix 1 contains 3 siRNAs (CliniSciences, CRH7929) and Mix 2 contains two siRNAs (ThermoFisher Scientific, 1299001 and 4392420). As a negative control, scramble siRNA was used (ThermoFisher Scientific, AM4611). Naïve H9 were transfected using the lipofectamine RNAiMAX transfection reagent (ThermoFisher Scientific, 13778030) according to the manufacturer’s recommendations. PXGL medium was changed 1 hour prior to transfection. After 24 hours, another round of transfection was performed. After 48h in total, cells were collected and RNA was extracted using the RNeasy Mini kit (Qiagen) according to the manufacturer’s recommendation.

### Raw data processing

RNA-seq, fastGRO-seq, CUT&RUN, and RAP-seq reads were trimmed using trim_galore (0.6.5) (https://github.com/FelixKrueger/TrimGalore) with a minimum length of 50 bp. Reads were then mapped to the human genome (hg38) and mouse genome (mm10) using bowtie2 (2.4.4) (Langmead and Salzberg, 2012) with the following parameters: --local --very-sensitive-local --no-unal --no-mixed --no-discordant --phred33 -L 10 -X 700. Reads were then deduplicated using MarkDuplicates from picard (2.23.5) (http://broadinstitute.github.io/picard/) with the following options: --CREATE_INDEX=true --VALIDATION_STRINGENCY=SILENT --REMOVE_DUPLICATES=true --ASSUME_SORTED=true. Bam files were sorted, filtered (minimum mapping quality=10) and indexed with samtools (1.13) (Danecek et al., 2021). Reads from mouse embryonic fibroblasts (MEFs) were discarded using XenofilteR package from R (4.1.1) (Kluin et al., 2018).

### RNA-seq and fastGRO-seq data analysis

For gene expression analysis, read counts were quantified for each gene with htseq-count from htseq (0.13.5) (Anders et al., 2015) using the following options: --stranded reverse -a 10 -t exon -i gene_id -m intersection-nonempty. The annotation file used: Homo_sapiens.GRCh38.90.gtf. Reads marked by special counters (no feature, ambiguous, too low aQual, not aligned, alignment not unique) were eliminated. For XIST isoforms reconstruction, Scallop (0.10.4) was used using the default parameters (Shao and Kingsford, 2017). XIST isoform quantification was performed using kallisto (0.46.2) on trimmed fastq with the option --rf-stranded (Bray et al., 2016). For that purpose, transcriptome fasta of exons/introns was generated from hg38 reference genome using BUSpaRse package under R (4.1.1), and was used to produce a kallisto index.

### Single cell expression data analysis

Fastq files from (Petropoulos et al., 2016) and (Zhou et al., 2019) were obtained from ENA (European Nucleotide Archive), under accession numbers PRJEB11202 and PRJNA431392, respectively. STAR (2.7.9a) was used for read alignment to GRCh38 reference human genome. STAR index was generated using the following options: STAR --runMode genomeGenerate --runThreadN 30 --genomeFastaFiles HS.GRCh38.fa -- limitGenomeGenerateRAM 168632718037 --outFileNamePrefix STAR_INDEX. --sjdbGTFfile Homo_sapiens.GRCh38.90.gtf. --genomeDir STAR_INDEX_petropoulos_2016 and -- sjdbOverhang 42 were used for Petropoulos data while --genomeDir STAR_INDEX and -- sjdbOverhang 149 were used for Zhou data.

For Petropoulos data: Reads (single-end) were aligned using the following options: STAR -- soloType SmartSeq --soloFeatures Gene --runThreadN 12 --genomeDir STAR_INDEX_petropoulos_2016 --readFilesCommand zcat --readFilesIn sample_A.fastq.gz --soloStrand Unstranded --soloUMIdedup NoDedup --outFileNamePrefix petro_2016/STAR_ALIGN_sample_A.STARSOLO. --outSAMtype BAM SortedByCoordinate --outSAMattrRGline ID:sample_A.

For Zhou data: Reads (paired-end) were aligned using the following options: STAR --soloType CB_UMI_Simple --soloCBwhitelist zhou_2019/whitelist.txt --soloFeatures Gene Velocyto -- runThreadN 5 --genomeDir STAR_INDEX --readFilesCommand zcat --readFilesIn multiplex_sample_A_R1.fastq.gz multiplex_sample_A_R2.fastq.gz --outFileNamePrefix zhou_2019/STAR_ALIGN_multiplex_sample_A.STARSOLO. --outSAMtype BAM SortedByCoordinate --outSAMattributes NH HI nM AS CR UR CB UB GX GN sS sQ sM -- soloCBstart 1 --soloCBlen 8 --soloUMIstart 9 --soloUMIlen 7 --soloCellFilter None -- soloBarcodeReadLength 0.

Finally, raw counts generated by STAR --soloFeatures Gene option were normalized as CPM (counts per million), log2-transformed and quantile normalized for subsequent analyses.

### SNP calling

We used a whole genome sequencing dataset of H9 hESCs to identify informative genomic SNPs along the X chromosome which can be used for allelic expression analysis (GSM1227088). Reads were aligned using bowtie2 and PCR duplicates were filtered out using MarkDuplicates from picard (2.22.0) as previously described. For proper tags formatting of BAM files, AddorReplaceReadGroups from picard was used with the following options: SO =“coordinate”, RGID=“id”, RGLB=“library”, RGPL=“platform”, RGPU=“machine”, RGSM=“sample”. BAM files were further processed using GATK (4.1.2.0) (McKenna et al., 2010) for SNP identification. BaseRecalibrator and ApplyBQSR were used to generate recalibrated BAM using high confidence SNPs referenced in Mills_and_1000G_gold_standard.indels.hg38.vcf.gz (annotation of indels) and dbsnp_146.hg38.vcf.gz (annotation of SNPs). Variant Call Format (VCF) file H9_WGS_hg38_filtered.vcf was produced with HaplotypeCaller (options: --dont-use-soft-clipped-bases and -stand-call-conf 10.0) and filtered using VariantFiltration (options: -window 50, -cluster 3, --filter-name FS, -filter ‘FS > 30.0’, --filter-name QD, -filter ‘QD < 2.0’). SNP position file info_snp_heterozygous_h9_wgs_chrX.txt was produced from VCF under R (4.1.1), using minimum threshold 50 for quality score, 10 for read coverage and 0.25 for allelic ratio to define heterozygous positions. We then used mpileup from samtools (1.15.1) to produce pileup files from RNA-seq and fastGRO-seq BAM files with the following options: -- output-MQ, --no-output-ins, --no-output-ins, --no-output-del, --no-output-del, --no-output-ends. Putative Xa haplotype was generated under R (4.1.1) from monoallelically expressed SNPs from the X chromosome in H9 primed hESCs. This was used to determine pXi/Xa allelic ratio across all samples. We considered a transcript as biallelic when at least 25% of reads originated from the second allele.

### CUT&RUN data analysis

BigWig track files were generated with bamCoverage from deeptools (3.5.0) using the following parameters: --normalizeUsing BPM --binSize 20 --smoothLength 40. Coverage score matrices were generated with computeMatrix from deeptools (3.5.0) using either reference-point --referencePoint TSS -R Homo_sapiens.GRCh38.90.gtf -- beforeRegionStartLength 1000 --afterRegionStartLength 1000 --binSize 20 -- missingDataAsZero or scale-regions -R Homo_sapiens.GRCh38.90.gtf --regionBodyLength 20000 --startLabel TSS --endLabel TES --binSize 20 –missingDataAsZero. For histone modification enrichment, raw counts were generated using multiBigwigSummary from deeptools with --binSize 10000 and the mean counts per chromosome was calculated. Percentage occupancy was determined by calculating for each chromosome the cumulative coverage over the chromosome size. Regions with a minimum threshold of 1.2 (fold change over IgG) are considered covered.

### ATAC-seq data analysis

ATAC-seq raw data were analyzed using the atacseq pipeline (1.2.1) (https://github.com/nf-core/atacseq/) from nf-core (Ewels et al., 2020) using the default parameters. Briefly, reads were aligned to the human genome (hg38) using BWA (0.7.17). Duplicates were marked by picard and reads mapping to mitochondrial DNA and blacklisted regions were removed. BigWig files were generated using deeptools as described above.

### MeD-seq data analysis

Data processing was carried out using specifically created scripts in Python. Raw fastq files were subjected to Illumina adaptor trimming and reads were filtered based on LpnPI restriction site occurrence between 13-17 bp from either 5’ or 3’ end of the read. Reads that passed the filter were mapped to hg38 using bowtie2. Genome-wide individual LpnPI site scores were used to generate read count scores for the following annotated regions: transcription start sites (TSS, 1 kb before and 1 kb after), CpG-islands and gene bodies (1kb after TSS till TES) and normalized (RPM) using total amount of CpG reads after filter. Gene and CpG-island annotations were downloaded from ENSEMBL (www.ensembl.org).

### RAP-seq data analysis

BigWig tracks were generated as previously described. RAP-seq naïve vs primed log2 enrichment was calculated using bigwigCompare from deeptools using the default parameters. Percentage occupancy was determined by calculating for each chromosome the cumulative coverage over the chromosome size. Regions with a minimum threshold of 10 (fold change over Input) are considered covered. RAP-seq coverage scores were calculated with multiBigwigSummary, using bins option for gene density enrichment and BED-file option for transposable elements (annotation from RepeatMasker).

### Quantification and statistical analysis

Statistical analyses were performed in R (4.1.1). The statistical analysis method for each experiment is specified in the figure legend. P-values <0.05 were considered statistically significant. Unless otherwise mentioned, most of the data shown are either representative of three or more independent experiments, or combined from three or more independent experiments as noted and analyzed as mean ± SD.

## Data availability

All RNA-seq, RAP-seq, ATAC-seq, CUT&RUN, MeD-seq, and fastGRO-seq data have been deposited at GEO (XXX) and are publicly available as of the date of publication. Microscopy data reported in this paper will be shared by the lead contact upon request. This paper analyzes existing, publicly available data. scRNA-seq data from human embryos are available at ENA (European Nucleotide Archive), under accession numbers PRJEB11202 and PRJNA431392. Hi-C data for H9 hESCs available at GEO GSE133126. CUT&RUN and Minute ChIP data for naïve H9 hESCs are available at GSE176175 and GSE181244, respectively.

## Code availability

All original code has been deposited at GitHub (URL) and is publicly available as of the date of publication. Any additional information required to reanalyze the data reported in this paper is available from the lead contact upon request.

