## Supplementary Figures for "XIST dampens X chromosome activity in a SPEN-dependent manner during early human development"

Supplementary Figure 1: Characterization of the X chromosome status in naive XIST KO, XIST CRISPRi, and XACT KO clones

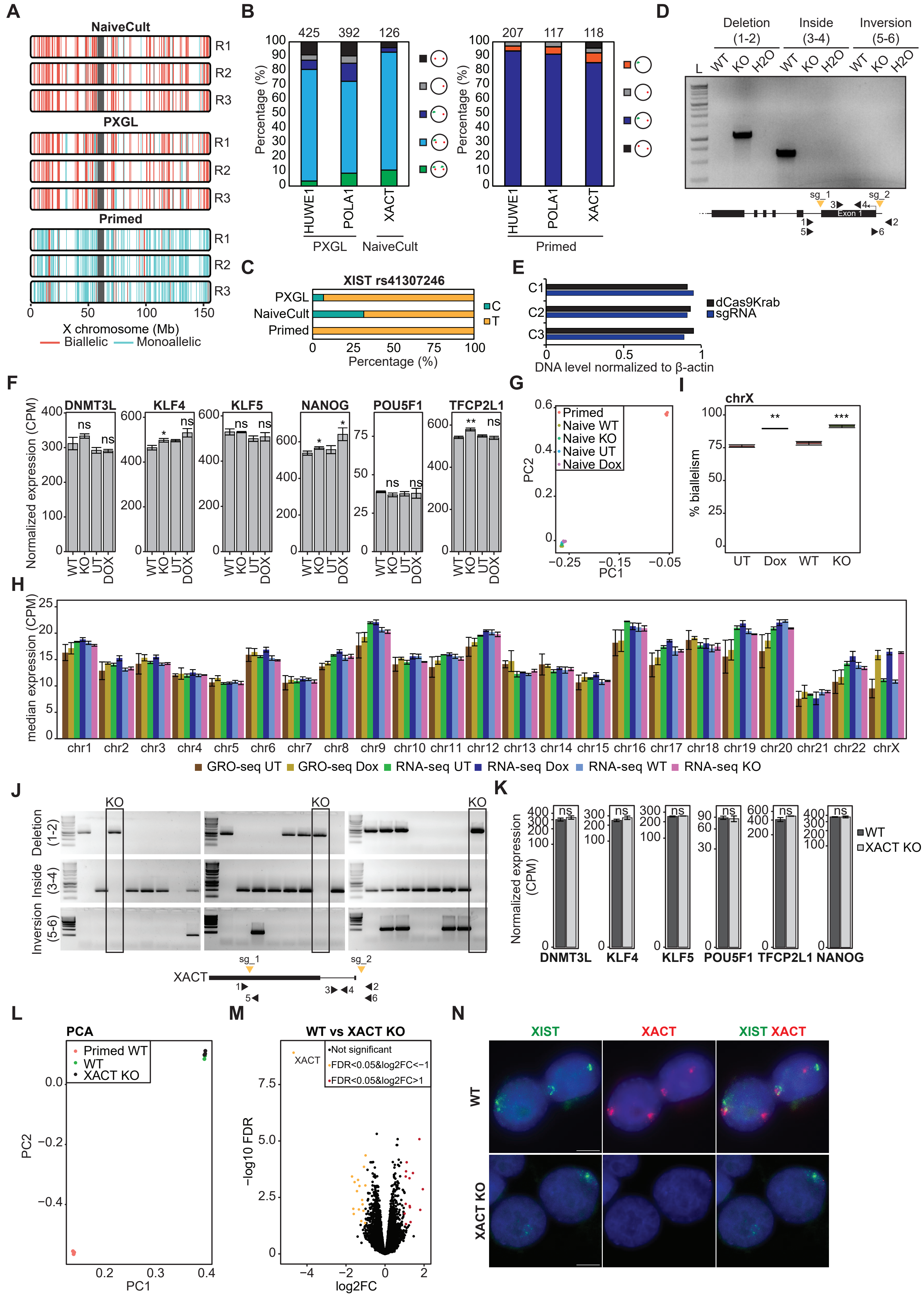

Supplementary Figure 2: Characterization of XIST enrichment on the X chromosome in naive and primed hESCs

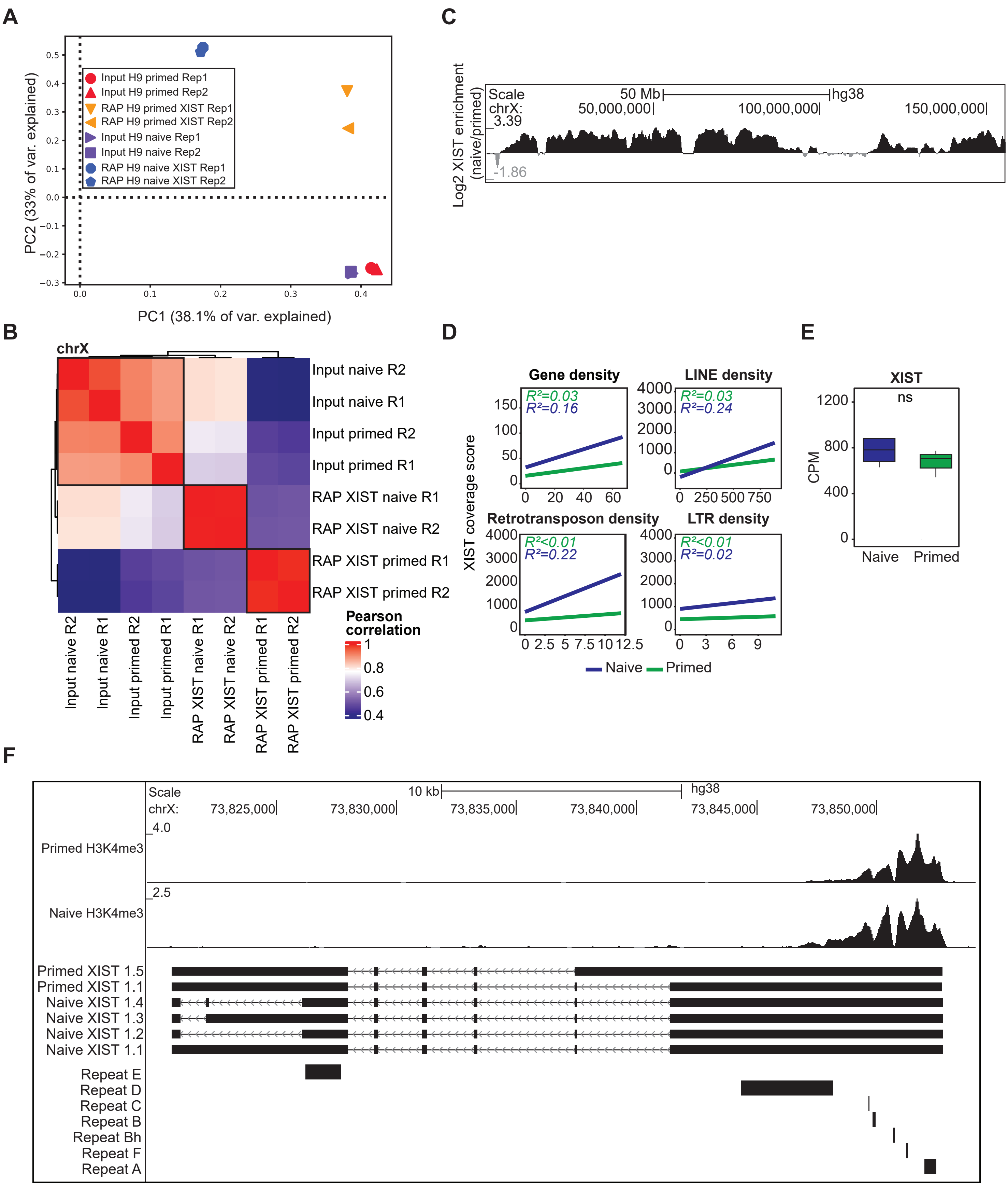

Supplementary Figure 3: Characterization of the X chromatin landscape in WT, XIST KO and XIST CRISPRi UT and Dox-treated clones in naïve hESCs

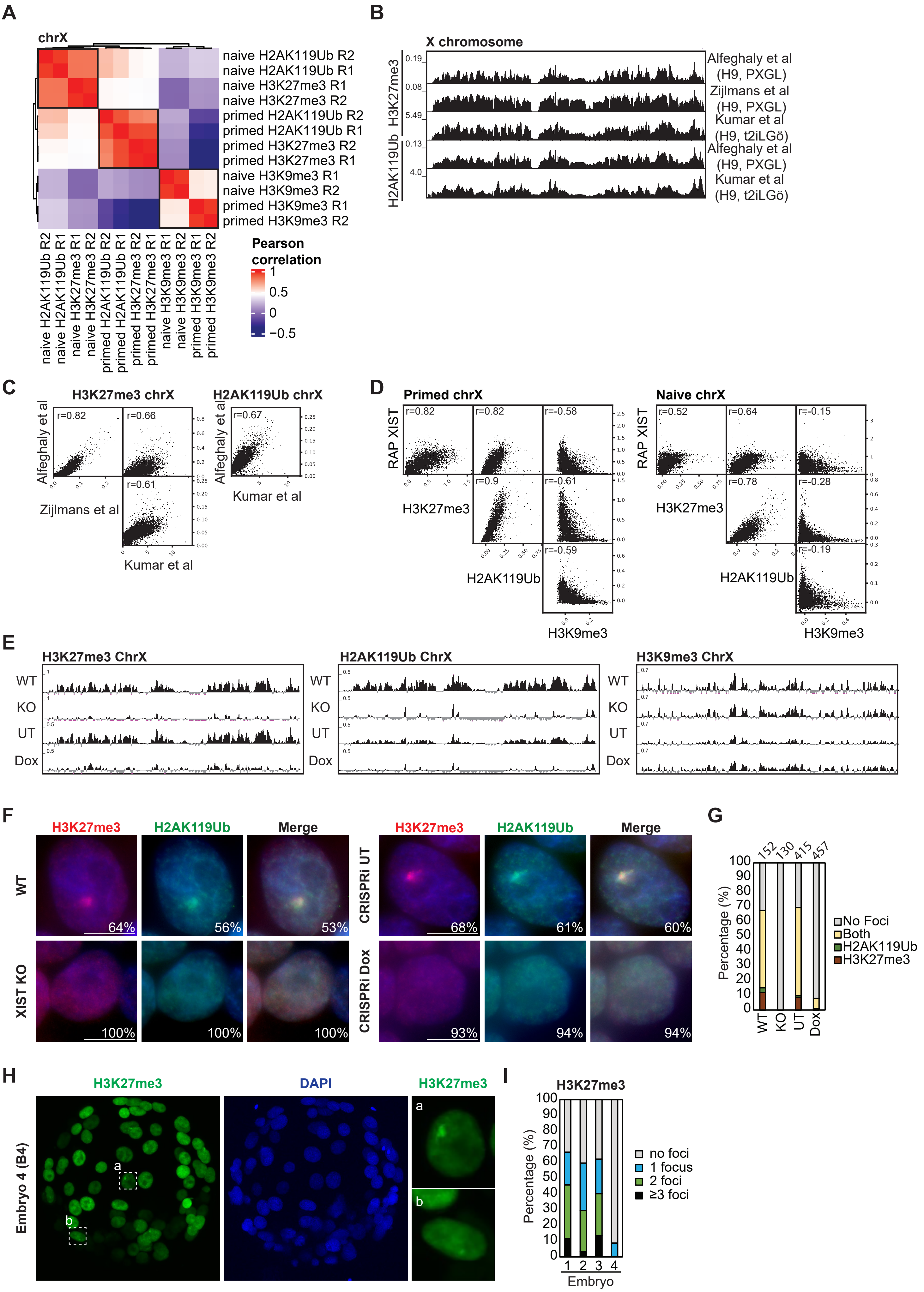

Supplementary Figure 4: Genetic and epigenetic features of XIST-sensitive and XIST-resistant genes

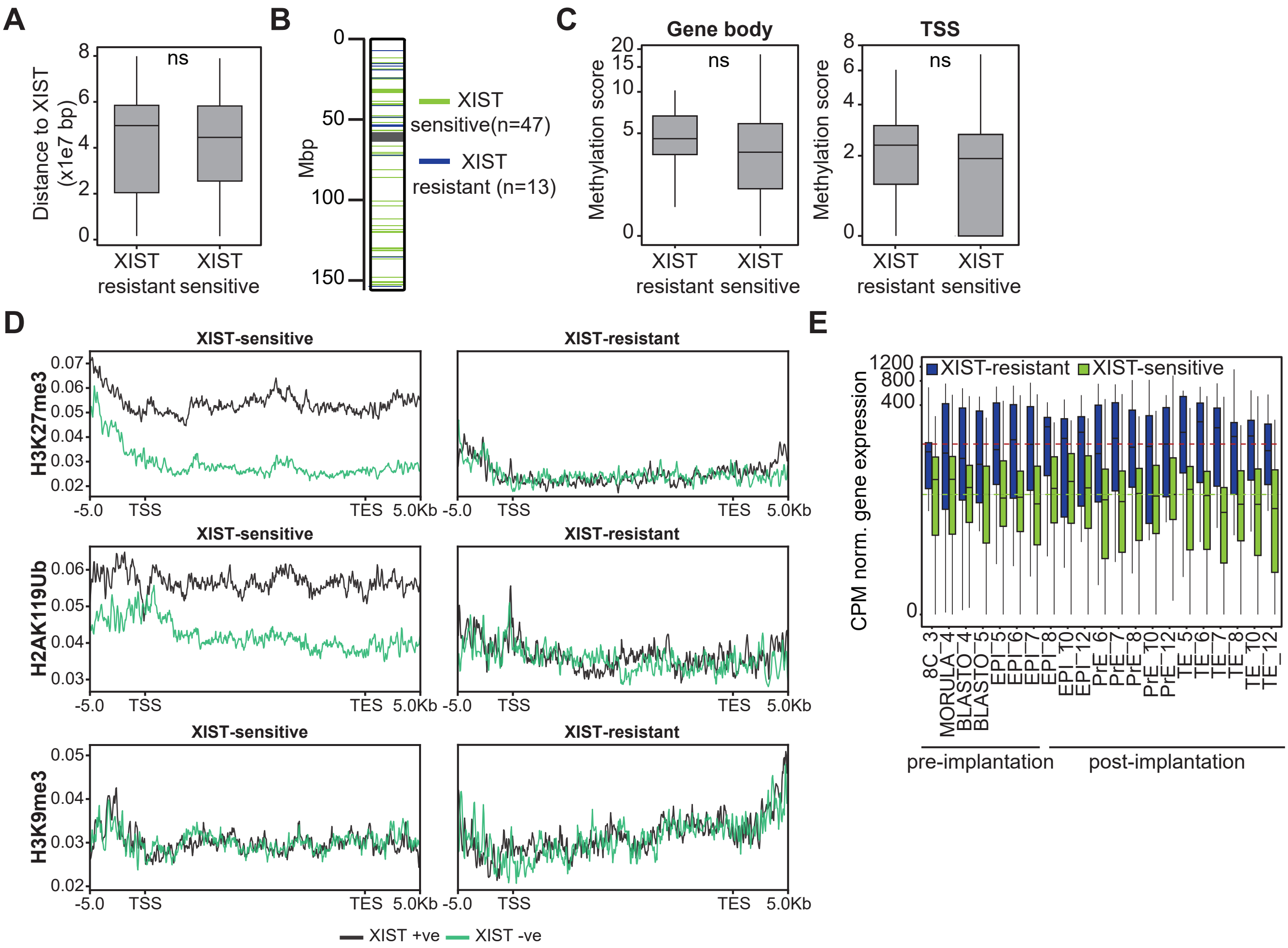

Supplementary Figure 5: Characterization of SPEN expression and KD

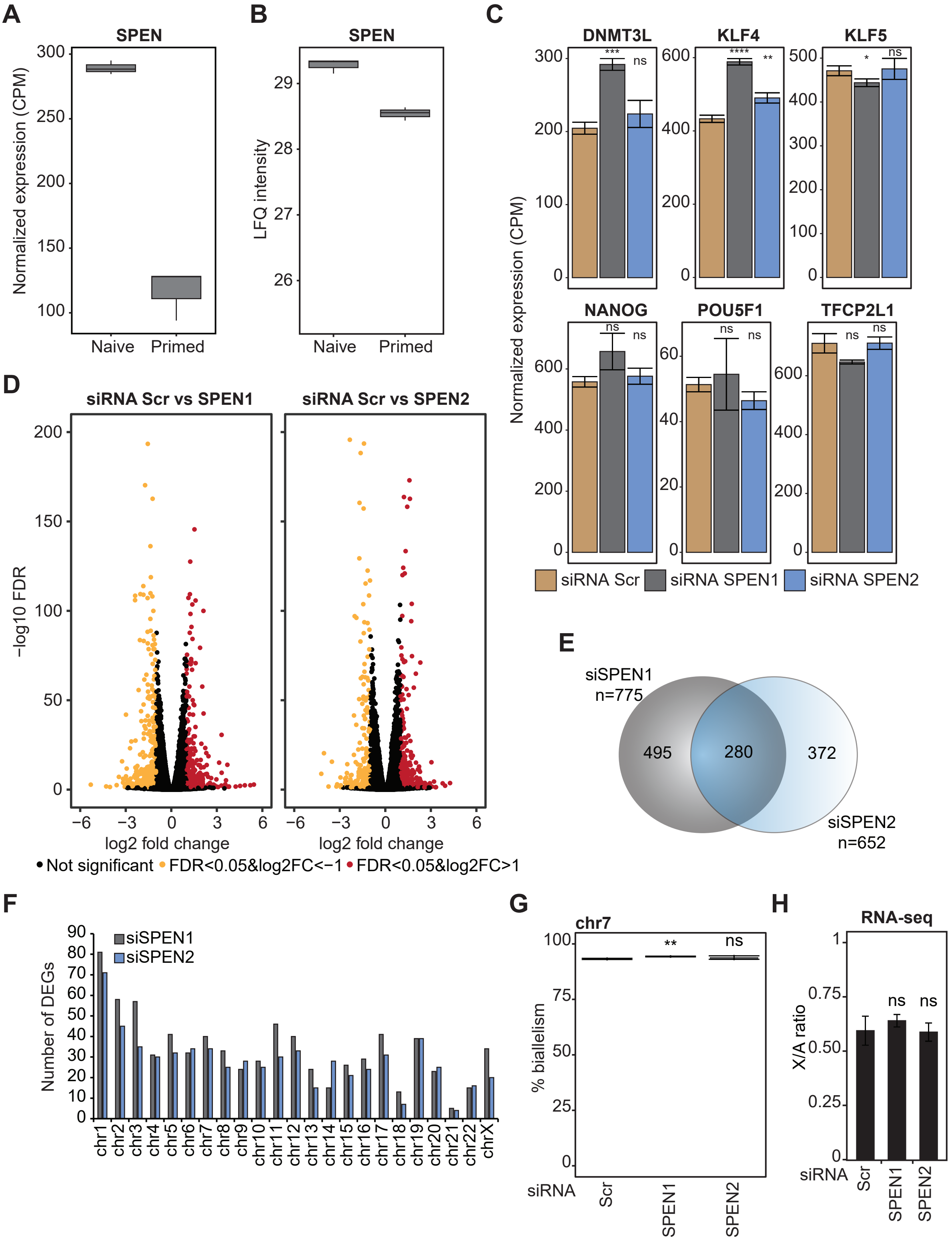
